# Status of Round Goby Invasion Fronts in New York and Quebec: Implications for Lake Champlain

**DOI:** 10.64898/2026.03.23.712452

**Authors:** Scott D. George, Hannah L. Diebboll, Steven H. Pearson, Jesica Goldsmit, Annick Drouin, Nathalie Vachon, Guillaume Côté, Siena Daudelin, Meredith L. Bartron, Meg D. Modley, Kate A. Littrell, Rodman G. Getchell, Rob J. Fiorentino, Thomas R. Sadekoski, Jason S. Finkelstein, Michael J. Darling, Geneviève J. Parent, Lauren M. Atkins

## Abstract

Invasive round goby *Neogobius melanostomus* have advanced eastward through the state of New York and provinces of Ontario and Quebec over the past two decades and are approaching Lake Champlain, one of the largest lakes in North America. This manuscript describes international efforts to monitor round goby populations during 2021–2025 on (a) the southern approach to Lake Champlain via the Hudson River and Champlain Canal, and (b) the northern approach to Lake Champlain via the Saint Lawrence River and Richelieu River. Monitoring utilized environmental DNA (eDNA), backpack electrofishing, beach seining, benthic trawling, and viral hemorrhagic septicemia virus (VHSV) testing. In the Champlain Canal, round goby were captured as far north as the downstream side of the C1 dam (97 kilometers [km] from Lake Champlain) while eDNA detections occurred as far north as the upstream side of the C2 dam (90 km from Lake Champlain). In the Richelieu River, round goby were captured as far south as Saint-Marc-sur-Richelieu (82 km from Lake Champlain) while the southern-most eDNA detections occurred near the Canadian side of the international border (4 km from Lake Champlain). Water temperature influenced habitat usage of round goby in the Champlain Canal, with catch rates in near-shore areas declining at < 10 °C. All VHSV test results were non-detections at the mouth of the Richelieu River, while one positive and two inconclusive results occurred along the Champlain Canal. Together, these data have informed multiple mitigation measures and have implications for management of aquatic invasive species across North America.

## Introduction

Since being first identified in North America in 1990, round goby *Neogobius melanostomus* have been found in all five Great Lakes, the Saint Lawrence River, and numerous inland waters. By the end of 2025, the species had been reported from two Canadian provinces and nine U.S. states (U.S. Geological Survey 2025), and inhabited watersheds covering at least 3 % of Canada and 11 % of the conterminous U.S. (HydroBASINS level 5 (Lehner and Grill 2013)). The rapid spread of this invasive species is attributed to its high reproductive potential and an unknown number of human-assisted movements (Kornis et al. 2012; Mueller et al. 2017; Johansson et al. 2018).

Establishment of round goby populations is associated with numerous ecological impacts including displacement of native benthic fish, egg predation of desirable gamefish, and transfer of contaminants and toxins to higher trophic levels (Corkum et al. 2004; Kornis et al. 2012). The species is also increasingly recognized as the most prevalent carrier of the viral hemorrhagic septicemia virus (VHSV) (Haws et al. 2025), which has caused fish kills in the Great Lakes region and is associated with declining muskellunge *Esox masquinongy* populations in the Saint Lawrence River (Farrell et al. 2017). However, round goby can also function as an abundant forage species (Reyjol et al. 2010), and have been associated with increased growth rate of some predators (Crane et al. 2015), while potentially also relaxing predation pressure on other prey species (Morissette et al. 2018).

Round goby have advanced eastward over the past two decades from the initial introduction in the Great Lakes into the State of New York and the province of Quebec (Canada). In New York (NY), round goby invaded east and south from their Lake Erie and Lake Ontario populations towards the interior of the state. They were identified in Onondaga Lake in 2010, Cayuga Lake in 2011, and Oneida Lake in 2013 where the species became the most abundant benthic fish by 2015 (Jackson et al. 2016). In 2014, a single round goby was captured by an angler in the Erie Canal approximately 40 kilometers (km) east of Oneida Lake near Utica, and in 2020 another angler-caught specimen occurred approximately 9 km upstream of the confluence of the Mohawk and Hudson Rivers (U.S. Geological Survey 2025). The first capture of round goby in the Hudson River occurred in 2021 near Albany (Figure 1) during a routine seining survey (Pendleton et al. 2022). The establishment of a round goby population in the Hudson River sparked considerable media attention and concern amongst natural resource managers and conservation groups that the species might advance upstream through the Hudson River and Champlain Canal, ultimately reaching Lake Champlain.

**Figure 1.**
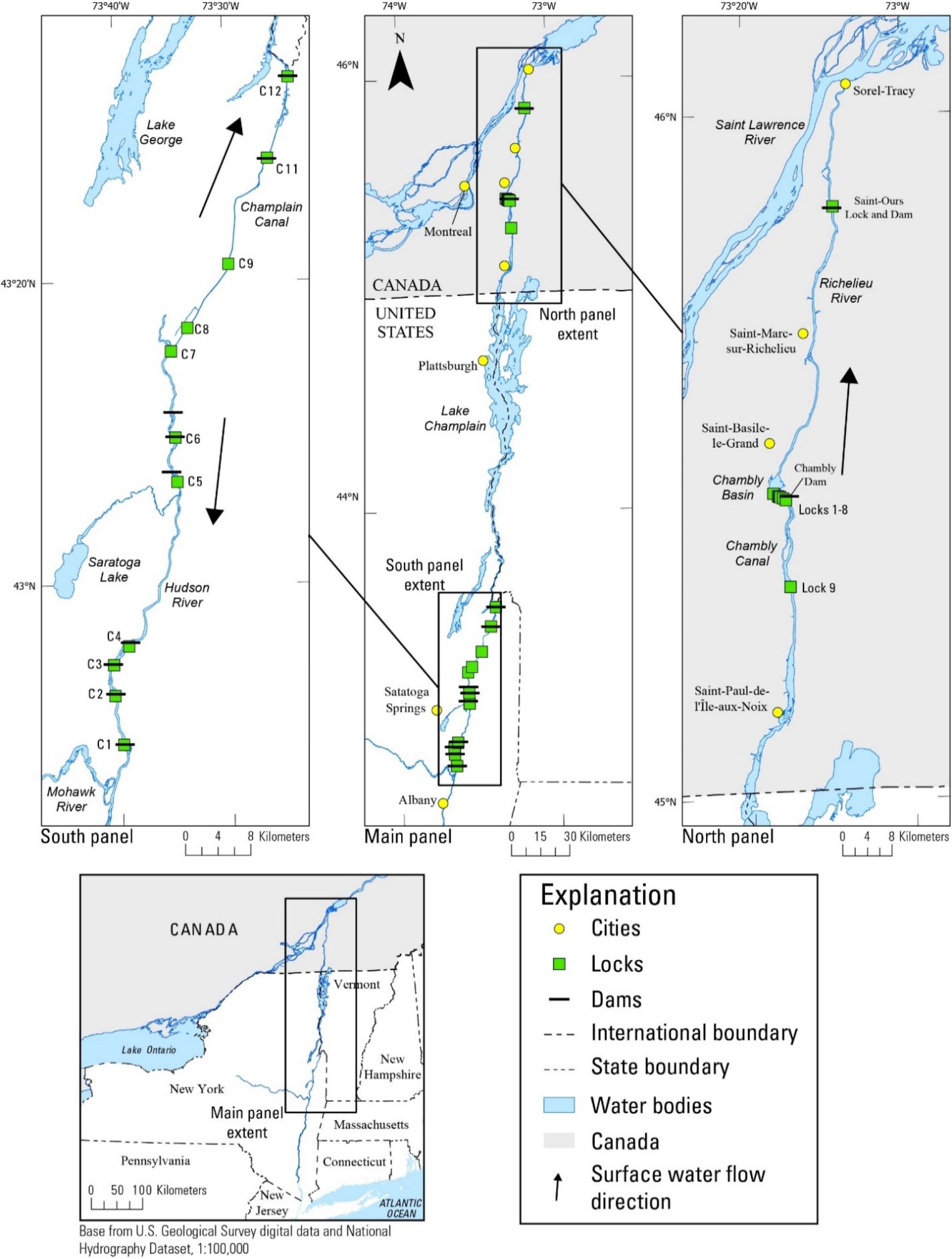
Map of study area with insets of Champlain Canal (left) and Richelieu River (right).

In Quebec, round goby were first identified in the Saint Lawrence River in 1997. Round goby were infrequently captured over the next decade but became abundant in most parts of the Saint Lawrence River from St. Francis Lake to Becancour-Batiscan between 2007 and 2011 (Morissette et al. 2018). Since then, round goby have expanded further, reaching the start of brackish waters in the Saint Lawrence River (MELCCFP 2025). Round goby were first captured in the Richelieu River in 2011 (Vachon 2018). At the initiation of this study in 2021, round goby were abundant between the mouth of the Richelieu River and Saint-Ours Dam (approximately 23 km upstream; Figure 1) but unreported upstream of the dam (MELCCFP 2025).

Lake Champlain is among the 40 largest lakes in North America, and its waters span parts of New York, Vermont, and Quebec. Its 21,326 km^2^ watershed is 73% forested but also includes fertile agricultural river valleys (LCBP 2024). Although the lake is known to have at least 51 aquatic non-native species (Marsden and Hauser 2009; LCBP 2024), numerous high-profile invasive species such as quagga mussel *Dreissena bugensis*, hydrilla *Hydrilla verticillata*, and northern snakehead *Channa argus* have not been reported. Lake Champlain is a high-profile destination for tournament bass angling and hosts national-circuit events, alongside numerous smaller tournaments each year, contributing to the regional economy. As a partial indicator of ecosystem-service benefits, Voigt et al. (2015) estimated that Lake Champlain-related tourism and second-home spending in four lakeshore Vermont counties in 2013 supported over $200 million USD in regional value added – indicating that a complete and contemporary lake-wide valuation would be considerably higher. Lake Champlain and its tributaries also support nine fish species and eight freshwater mussel species listed as threatened or endangered in New York, Vermont, or Quebec (LCBP 2022). Additionally, lake trout *Salvelinus namaycush*, a native gamefish that was formerly extirpated from the lake, has recently been recognized as a self-sustaining population that no longer requires supplemental stocking (USFWS 2025). As a result, environmental stressors and invasive species that threaten this ecosystem are of great concern to local communities, natural resource managers, and elected officials.

Most information indicates Lake Champlain is suitable, and potentially excellent, habitat for round goby. Its waters are characterized by rocky shorelines and hard substrates that are heavily colonized by invasive zebra mussel *Dreissena polymorpha*, an important food source for round goby in both its native and non-native ranges. Additionally, the lake spans a range of trophic conditions and has a mean depth of 20 meters (m) (Cohn et al. 2007) – both features that should be favorable to colonization. The threat of round goby invasion is especially concerning given the well-established role of round goby as an egg predator, and the affinity of the species for similar habitats as those used for lake trout spawning. These characteristics have raised concerns as to whether round goby could reverse decades of progress on lake trout restoration. Lake Champlain also supports a small population of muskellunge which could be imperiled by the establishment of round goby and its role as a disease reservoir for VHSV (Farrell et al. 2017; Haws et al. 2025).

The primary objective of the multi-agency effort described herein was to collect fine-scale spatial and temporal information on the distribution of round goby in connecting waters to Lake Champlain. Additional goals of this research were to (a) determine if the invasion front was carrying VHSV, (b) estimate rates of round goby expansion, and (c) collect baseline information on existing fish communities that could be used to quantify future change. Examples of how these data are being used within management plans to inform operations of infrastructure and adaptive sampling efforts are also described.

## Methods

This research was conducted in two geographic areas representing the southern and northern approaches to Lake Champlain. Monitoring on the southern approach was conducted on a 60-km section of the Champlain Canal between Waterford and Fort Edward, NY. This section is a canalized part of the upper Hudson River in which water levels are seasonally maintained for navigation. It contains seven locks (C1-C7) and seven dams (most of which are co-located) (Figure 1). The downstream-most dam at lock C1 has six tainter gates, each blocking 8.8-m tall and 15.2-m wide gate bays. The tainter gates are lifted during winter and flow through the gate bays drains the navigation pool to protect infrastructure from ice damage. The remaining six dams upstream of C1 are fixed and do not have tainter gates. The flow direction in this section is from north to south, meaning the movement of species towards Lake Champlain occurs in an upstream direction.

Research on the northern approach to Lake Champlain was conducted on the 124-km Richelieu River. The Richelieu River is the outlet of Lake Champlain and flows north until reaching the Saint Lawrence River in Sorel-Tracy (Figure 1). The Richelieu River has two major dams. Moving upstream from the Saint Lawrence River, the first is the Saint-Ours Dam at river kilometer 23. The dam has an adjacent lock that is operated seasonally from May to October as well as a fishway that enables upstream passage of many fish species. The second is the Chambly Dam which occurs at river kilometer 75 and is impassable for upstream movement of fish, except for American eel *Anguilla rostrata* via an eel ladder. The 19-km Chambly Canal National Historic Site allows for seasonal boat navigation around the Chambly Dam and Chambly Rapids and is composed of nine locks.

The methods and sampling regimes used for monitoring the southern and northern approaches had broad similarities in their structure and objectives but also substantial methodological differences as a result of leveraging preexisting monitoring programs and the complexities of coordinating international, multi-agency efforts. Major similarities included the combined use of environmental DNA (eDNA) and physical capture methods in each area. For eDNA, all agencies filtered 2 liters (L) of water (or until clogging) through a 47 millimeter (mm) filter with a pump system, used distilled or deionized water for negative field controls, extracted DNA using the Lyse and Spin basket and DNeasy Blood and Tissue kit (Qiagen, Germantown, MD, USA), and detected target DNA with a quantitative polymerase chain reaction (qPCR) based assay. Samples were classified as positive if at least one qPCR replicate amplified for the target sequence within the prescribed number of cycles. This sensitive detection criterion was applied to minimize false negatives and maximize detection probability given the low environmental DNA concentrations expected near the invasion front. Despite these similarities, the comprehensive effort described herein involved eDNA sample collection by four agencies, eDNA sample analysis by three laboratories, and physical fish collections by three agencies. The key differences and details unique to each monitoring program are described below.

### Champlain Canal sampling

In the Champlain Canal, monitoring relied on three sampling techniques: eDNA, backpack electrofishing, and benthic trawling. Environmental DNA samples were collected (a) by the U.S. Geological Survey (USGS) at seven fixed sites, four times annually (April, June, August, October) from 2022 through 2025 (n samples = 116), and (b) by the New York State Department of Environmental Conservation (NYSDEC) at 3-9 sites monthly (excluding winter) from July 2024 through November 2025 (n samples = 115). All samples were collected just below the water surface in near-shore habitats. Samples collected by USGS were taken using a handpump to vacuum water through 1.5-micron (μm) nominal pore-size glass-fiber filters. A field control was collected before sample collection at the first and last site sampled each day. The full summary of collection protocols and decontamination procedures are described in George et al. (2022). Samples collected by NYSDEC were taken using the Smith-Root eDNA backpack sampler (Vancouver, WA, USA) with Smith-Root self-preserving (desiccating) 5-μm polyethersulfone (PES) filters following USFWS (2024). A single field control was taken during each sampling event.

All eDNA samples collected from the Champlain Canal were analyzed by the U.S. Fish and Wildlife Service Northeast Fishery Center. DNA extraction and qPCR methods generally followed procedures for filter samples described in USFWS (2019). Eight qPCR replicates were analyzed for each sample using the ReesCOI molecular marker (George et al. 2021). Quantification cycle (Cq) values less than or equal to 40 were considered positive detections for round goby DNA while values above 40 were considered non-detections.

Prior to qPCR analysis with the ReesCOI marker, all extracts were screened for PCR inhibition using the TaqMan Exogenous Internal Positive Control (IPC) Kit (Applied Biosystems, Waltham, MA, USA). Three replicate 20-microliter (μL) IPC reactions were run per sample, each containing 17 μL master mix (Environmental Master Mix 2.0, primer, probe, water) and 3 μL DNA template. Plates also included triplicate no-template controls (3 μL water) to establish reference Cq values for inhibition assessment. Samples exhibiting IPC Cq delay (ΔCq) > 1 relative to the mean of the no-template controls were treated with a Zymo OneStep Inhibitor Removal column according to manufacturer instructions (Zymo Research, Irvine, CA, USA). The full summary of laboratory equipment, reagents, and reaction conditions are summarized in George et al. (2022), and details on the ReesCOI qPCR marker can be found in George et al. (2021).

Backpack electrofishing was conducted twice annually (June and August) at the same seven fixed sites from 2022 through 2025, and on an approximately bi-weekly basis at a separate group of seven sites clustered upstream and downstream of the C1 dam from late 2022 through 2025. All electrofishing surveys were conducted in wadeable (< 1 m), near-shore habitats in a downstream-to-upstream direction for exactly 900 seconds of “on time.” Surveys were conducted with one person operating a Smith-Root Apex backpack electrofisher (USGS surveys) or a Halltech HT-2000 (Halltech Aquatic Research Inc., Guelph, Ontario, CAN) (NYSDEC surveys) and two to three people netting fish. Most round goby captured with electrofishing were measured in the field for total length to the nearest mm.

Benthic trawling was conducted twice annually (June and August) from 2022 through 2025 at the seven fixed sites. Trawling was conducted using a net from Innovative Net Systems (Milton, LA, USA) similar to a mini-Missouri trawl and followed methods suggested by Herzog et al., (2009) and described in George et al., (2021). The trawl was deployed from the bow of a 17-foot (5.2 m) skiff with a 60 hp outboard motor and pulled in a downstream direction (operating the engine in reverse) for 150 m at approximately 2.5-3.0 km/hour. Three trawl pulls (replicates) were conducted at each site; one in the center of the river channel and one along the shoulders of each bank. The sampled depths generally ranged from 1.2 to 3.7 m along the channel shoulders and from 3.4 to 5.2 m in the center of the channel. All round goby captured with trawling were measured for total length and weight.

Two batches of round goby were collected annually for VHSV testing from 2022 through 2025 using backpack electrofishing. Each year, one batch was collected in May and another in June to maximize detection potential (Eckerlin et al. 2011; Cornwell et al. 2014). These collections were intended to sample the nearest location to the invasion front that had sufficiently high round goby density such that at least 20 individuals with a minimum total length of 50 mm could be captured. A location with sufficient round goby density was selected on the Mohawk River approximately 1.1 km upstream of its confluence with the Hudson River and the southern end of the Champlain Canal (Figure 2). There are no barriers in this 1.1 km stretch of river; therefore round goby collected from this site are considered representative of the broader population in the Champlain Canal. All batches were frozen at -20 °C and analyzed at the Cornell University Aquatic Animal Health Lab within 1-3 months of collection.

**Figure 2.**
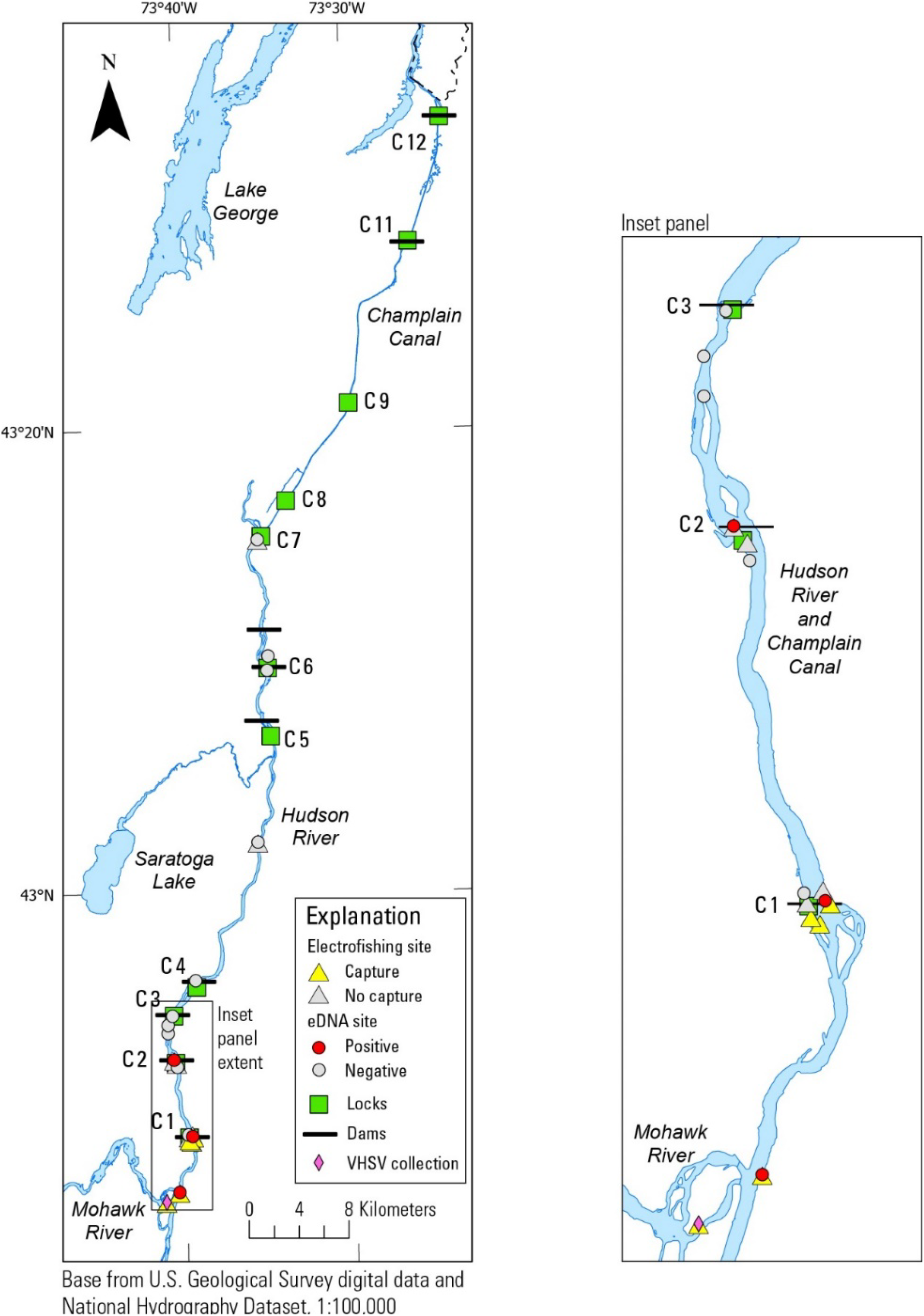
Environmental DNA (eDNA) and electrofishing results from round goby (Neogobius melanostomus) surveys on the Champlain Canal from 2022-2025. Data from some adjacent sites are combined for visual clarity.

A brain sample and a composite organ (spleen, heart, and kidney) sample from each specimen were analyzed in duplicate (two technical replicates) using quantitative reverse transcriptase polymerase chain reaction (qRT-PCR) for VHSV IVb as described in Hope et al. (2010), Standish et al. (2020), and George et al. (2022). Brain and composite organ samples were separately assigned a result of “non-detection”, “inconclusive”, or “positive”. A sample was classified as “positive” if the amplification curve for at least one replicate surpassed the detection threshold based on the standard curve, and the result was reproduced in both replicates during a subsequent rerun of the sample extract. A sample was classified as “inconclusive” if it amplified during the initial run and the result was not reproduced in both qRT-PCR replicates during a rerun of the sample extract.

### Richelieu River sampling

In the Richelieu River, monitoring was conducted with eDNA and beach seining. Environmental DNA samples were collected (a) by the Quebec Ministère de l’Environnement, de la Lutte contre les Changements Climatiques, de la Faune et des Parcs (MELCCFP) twice annually between July and September from 2021 through 2025 at sites that varied in location over time along the Richelieu River (n samples = 581), and (b) by Parks Canada three times annually (April, August, October) from 2023 through 2025 at 4-6 sites in the Chambly Canal (variable site locations; two field replicates per site; n samples = 99). The MELCCFP collected depth-integrated water samples by boat from water depths ranging from 0.5 to 10 m, while Parks Canada collected water samples by wading from immediately below the surface (2023-2024) or just above the bottom (2025) in depths up to 3.5 m. All water samples were filtered using a Smith-Root eDNA Citizen Scientist sampler to suction water through a Smith-Root self-preserving (desiccating) 1.2-μm pore size PES filter (MELCCFP and Parks Canada) or 1.5-μm nominal pore size glass-fiber filter (Parks Canada). Glass-fiber filters were preserved with silica beads. A field control was collected for every 14 samples during the MELCCFP surveys and after the last site sampled each day during the Parks Canada surveys. All field sampling followed contamination-prevention procedures described in Chevrinais et al. (2023).

All MELCCFP and Parks Canada samples were analyzed with six qPCR replicates using a new round goby qPCR assay (development and performance described in Supplemental Material). Values of Cq less than or equal to 50 were considered positive detections for round goby DNA, whereas no Cq values were considered non-detections. The MELCCFP samples were analyzed by the MELCCFP Complexe Scientifique eDNA laboratory as follows. Whole filter DNA extractions were performed under an ultraviolet (UV) hood with bleached and UV-treated instruments (Goldberg et al. 2011; Spens et al. 2017).

An extraction negative control was included in each extraction batch. Amplification was performed using the QuantStudio 5 qPCR platform (Thermo Fisher Scientific, Waltham, MA, USA) in a final volume of 20 μL including 1.8 μL of each primer (10 micromolar [μM]), 0.5 μL of probe (10 μM), 10 μL of Environmental Master Mix 2.0 (Thermo Fisher Scientific), 3.9 μL of SPUD (Nolan et al. 2006) and 2 μL of DNA extract under the following conditions: 2 minutes (min) at 50 °C, 10 min at 95 °C, 50 cycles of 15 seconds (s) at 95 °C, and 60 s at 60 °C. Each qPCR plate included six no-template controls (NTCs) and three positive controls (diluted round goby DNA). The SPUD assay served as an internal positive control for detecting inhibition. For each qPCR replicate, ΔCq ≥ 3 relative to the mean Cq of six plate-specific NTCs indicated PCR inhibition.

The Parks Canada samples were analyzed at the Maurice Lamontagne Institute of Fisheries and Oceans Canada as follows. DNA was extracted from a half filter and eluted in 100 μl Tris-buffer in an ultraclean room (Chevrinais et al. 2023). An extraction negative control composed of a half filter immersed in Milli-Q water (MilliporeSigma, Burlington, MA, USA) was included with the samples for each extraction batch. The qPCR reaction was run on a QuantStudio 3 (Thermo Fisher Scientific, MA, USA) in 20 μL of total volume, including 3 μL of extract, 0.4 μL NEME_F primer (10 μM), 1.2 μL NEME_F primer (10 μM), 0.4 μL of probe (10 μM), 10 μL of TaqPath™ ProAmp™ Master Mix 2X (Thermo Fisher Scientific, MA, USA), and completed with PCR grade water, under the following conditions: 10 min at 95 °C, 50 cycles of 15 s at 95 °C, and 60 s at 60 °C. Each qPCR plate included two NTCs and six positive controls, i.e., 2^*^10 copies and 4^*^2 copies per reaction of a quantified plasmid DNA (pDNA, Azenta Life Sciences, Burlington, MA, USA). Inhibition was tested with an IPC of 50 plasmid DNA copies (including the Frankenfish sequence, a chimera 12S sequence from three fish species (Chevrinais et al. 2023) added to a single qPCR replicate for each DNA extract, where a ΔCq ≥ 2 indicated PCR inhibition according to amplification conditions described in Chevrinais et al. (2023).

Physical captures of round goby in the Richelieu River primarily occurred as bycatch of a beach seining program to monitor young-of-the-year copper redhorse (*Moxostoma hubbsi*). This monitoring program has been taking place at a standardized group of 40 sites in Saint-Marc-sur-Richelieu in mid-to-late September every 1-2 years since 1997, and at additional sites downstream of the Saint-Ours Dam in some years (Vachon 2002; Vachon 2018; Vachon 2021). Additional targeted beach seining to monitor round goby distribution was performed in 2023 and 2024 upstream of Saint-Ours. All seining was conducted using a net of 4-m height, 12.5-m length, and a 3-mm stretched mesh size that is equipped with a fork with 12.5-m head and tail ropes. The seine was deployed with a boat and pulled over a standardized distance of 10 m toward the shore (depth of 0 m) for a sampled area of approximately 120 m^2^. Round goby captured with seining were preserved in 95% ethanol and then measured for total length.

Beach seining was used to capture round goby at the confluence of the Richelieu River and Saint Lawrence River for VHSV testing during May 2024. Round goby were sacrificed using a solution of 10% eugenol and subsequently stored at -20 °C. Testing for VHSV was performed at the Ontario Veterinary College. Tissues from the heart, gill, spleen and kidney from batches of five fish were composited in RNAlater (Thermo Fisher Scientific, Waltham, MA, USA) and frozen at -80 °C. Composited tissue samples were then homogenized, extracted using a total RNA kit, and analyzed in duplicate using qRT-PCR as originally described by Hope et al. (2010). Modifications included a standard curve generated from a synthetic DNA fragment (gBlock; Integrated DNA Technologies, Coralville, IA, USA) representing the VHSV target sequence.

Data collected by the USGS are publicly available in George et al. (2022; 2025). Data collected by the NYSDEC, MELCCFP, and Parks Canada are available upon request.

## Results

### Champlain Canal

All quality assurance data associated with eDNA samples were acceptable. No field controls detected the presence of round goby DNA, and all laboratory extraction and PCR controls produced expected results. Evidence of inhibition was only detected in four eDNA samples and was relieved as described above prior to qPCR analysis with the ReesCOI marker. Together, this information indicates that data quality was high, and the resulting data can be considered representative of ambient conditions in the environment.

A total of 231 eDNA samples were analyzed from the study area between 2022 and 2025. Thirty (13%) samples were classified as positive while 201 (87 %) were non-detections. Positive samples occurred almost exclusively at sites downstream of the Lock C1 dam and 88% of samples from that area were classified as positive (Figure 2). Two eDNA samples from all areas upstream of the Lock C1 dam were classified as positive, each occurring at the HR-263.0 site (one in June 2024 [5 of 8 qPCR replicates positive] and the other in August 2024 [1 of 8 qPCR replicates positive]). All samples subsequently collected from this location produced non-detections.

Electrofishing and trawling surveys collected a total of 633 round goby between 2022 and 2025. Captures occurred exclusively at sites located downstream of the Lock C1 Dam (Figure 2). Of note, an electrofishing survey in June 2022 captured 3 round goby immediately downstream of the C1 dam along the east shore. These specimens were the first documentation of upstream expansion from the confluence of the Mohawk and Hudson Rivers.

Of the 503 fish for which length was measured, the mean length was 50.4 mm (range 23-111 mm, n=480) from electrofishing captures and 31.0 mm (range 12-82 mm, n=23) from trawling captures. The 12 smallest round goby were captured via trawling while the 15 largest round goby were captured via electrofishing (Figure 3).

**Figure 3.**
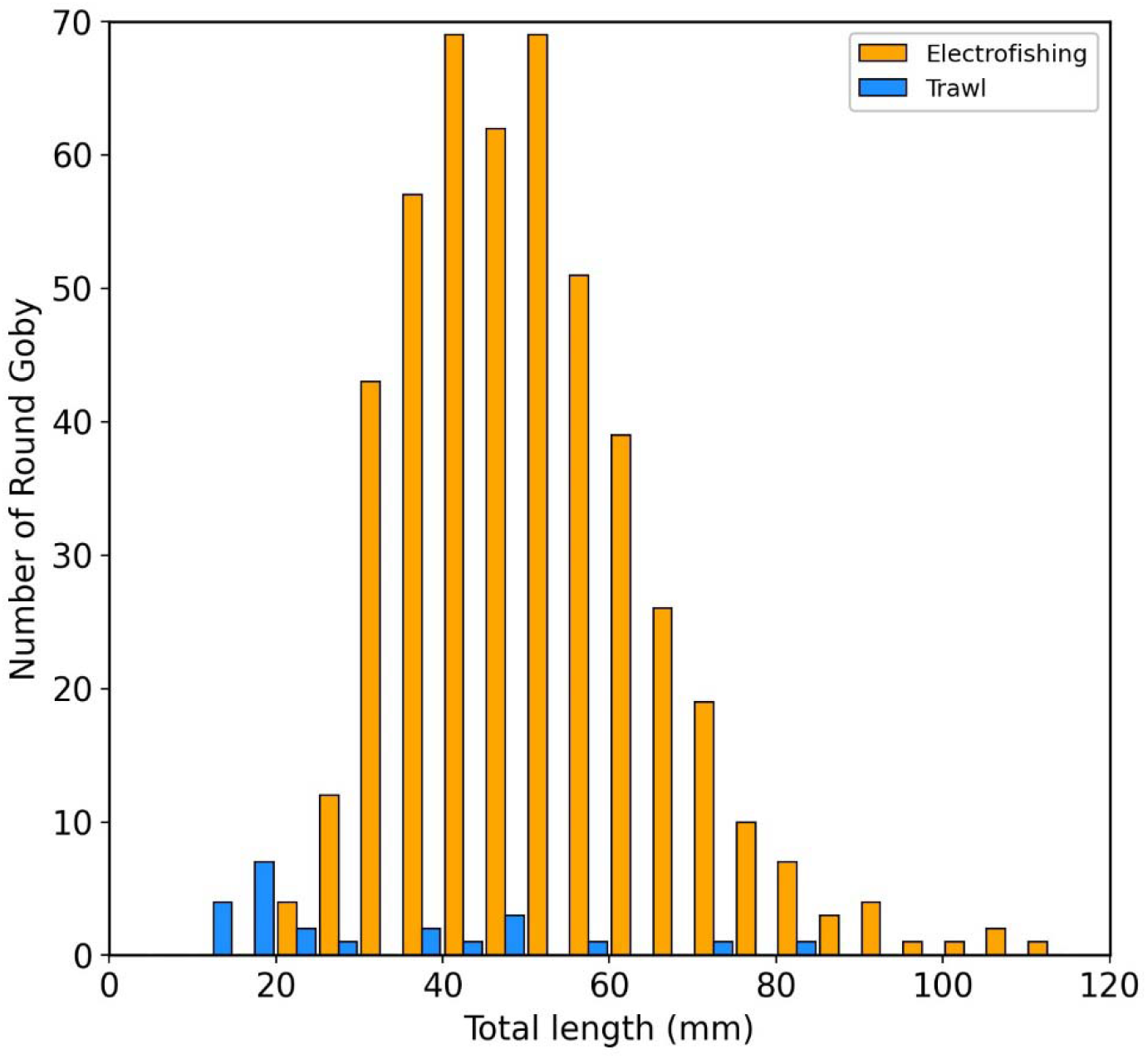
Total length-frequency distribution of round goby (Neogobius melanostomus) captured with routine monitoring at two sites between 2022 and 2025.

Water temperature appeared to influence the catch rate of round goby during electrofishing surveys. Data from trawling surveys were not considered in this analysis because the sampling was conducted less frequently than electrofishing and exclusively during warm-water periods. When data were aggregated from all electrofishing surveys conducted at sites downstream of the C1 dam (for which paired water temperature data were available), 84% of the round goby catch from the four-year study occurred at >10 °C. The mean catch per survey of round goby at the Hudson River downstream of Lock C1 site (HR-256.4) was 0.2 at ≤10 °C (n=13) and 1.0 at >10 °C (n=33) (Figure 4). Similarly, at the Mohawk River at Peebles Island site (MR-1.1), the mean catch was 6.5 at ≤10 °C (n=13) and 12.7 at >10 °C (n=33). Prior to November 2025, only 2 round goby had been captured across all surveys conducted at ≤ 5 °C and the overall dataset provided strong support for a water temperature effect on catch rate. However, the three most recent surveys conducted at the Mohawk River at Peebles Island site (November 19, 2025: 20 round goby at 5.0 °C, November 25, 2025, 40 round goby at 4.5 °C, and December 9, 2025, 11 round goby at 0.1 °C) strongly deviated from this pattern.

**Figure 4.**
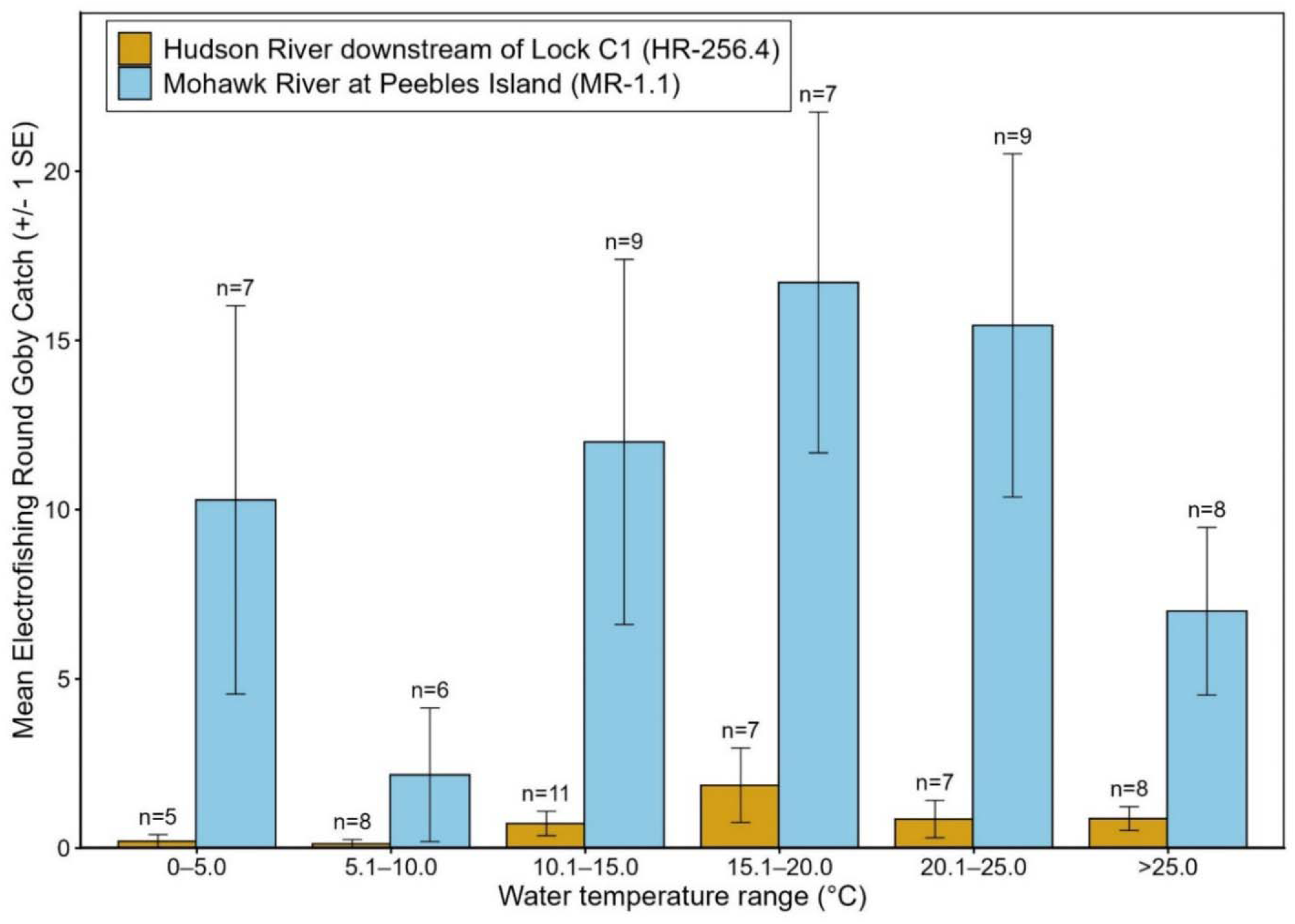
Catch (mean ± 1 standard error [SE]) of round goby (Neogobius melanostomus) from 900 second electrofishing surveys at two sites, binned by water temperature. The number of surveys represented by each bar is denoted by “n=“.

Two separate batches of round goby were tested for VHSV annually from 2022-2025. Of the 250 specimens that were tested, 0, 2, and 248 brain samples were classified as positive, inconclusive, and non-detection, respectively (Table 1). Similarly, 1, 0, and 249 composite organ samples were classified as positive, inconclusive, and non-detection, respectively.

**Table 1.** VHSV test results from batches of round goby collected near the Champlain Canal.

| Collection Period | Number of Round Goby | Brain Samples<br>(Positive/Inconclusive/Non-detection) | Organ Samples<br>(Positive/Inconclusive/Non-detection) |
| --- | --- | --- | --- |
| May 2022 | 34 | 0/0/34 | 0/0/34 |
| June 2022 | 36 | 0/1/35 | 0/0/36 |
| May 2023 | 30 | 0/0/30 | 0/0/30 |
| June 2023 | 30 | 0/0/30 | 0/0/30 |
| May 2024 | 30 | 0/0/30 | 0/0/30 |
| June 2024 | 30 | 0/0/30 | 0/0/30 |
| May 2025 | 30 | 0/0/30 | 0/0/30 |
| June 2025 | 30 | 0/1/29 | 1/0/29 |

### Richelieu River

The new eDNA qPCR assay for round goby was specific and sensitive (Supplemental Material). No cross amplification was observed for the 32 native species tested. A limit of detection (LOD) of 0.3 copies/reaction and a limit of quantification (LOQ) of 1.0 copies/reaction were determined with the qPCR conditions from the Maurice Lamontagne Institute (Supplemental Material). The MELCCFP did not determine the LOD or LOQ for the assay in their laboratory, as any amplification observed in a qPCR replicate prior 50 cycles was considered as a positive result.

All associated eDNA quality assurance data were acceptable. Round goby DNA was not detected in any of the field controls, no evidence of inhibition was detected, and all laboratory extraction and PCR controls produced expected results.

A total of 680 eDNA samples from the Richelieu River were analyzed for round goby DNA between 2021 and 2025. Positive samples accounted for 4.3 % of the sampling effort and occurred in a scattered pattern, both temporally and spatially (Figure 5).

**Figure 5.**
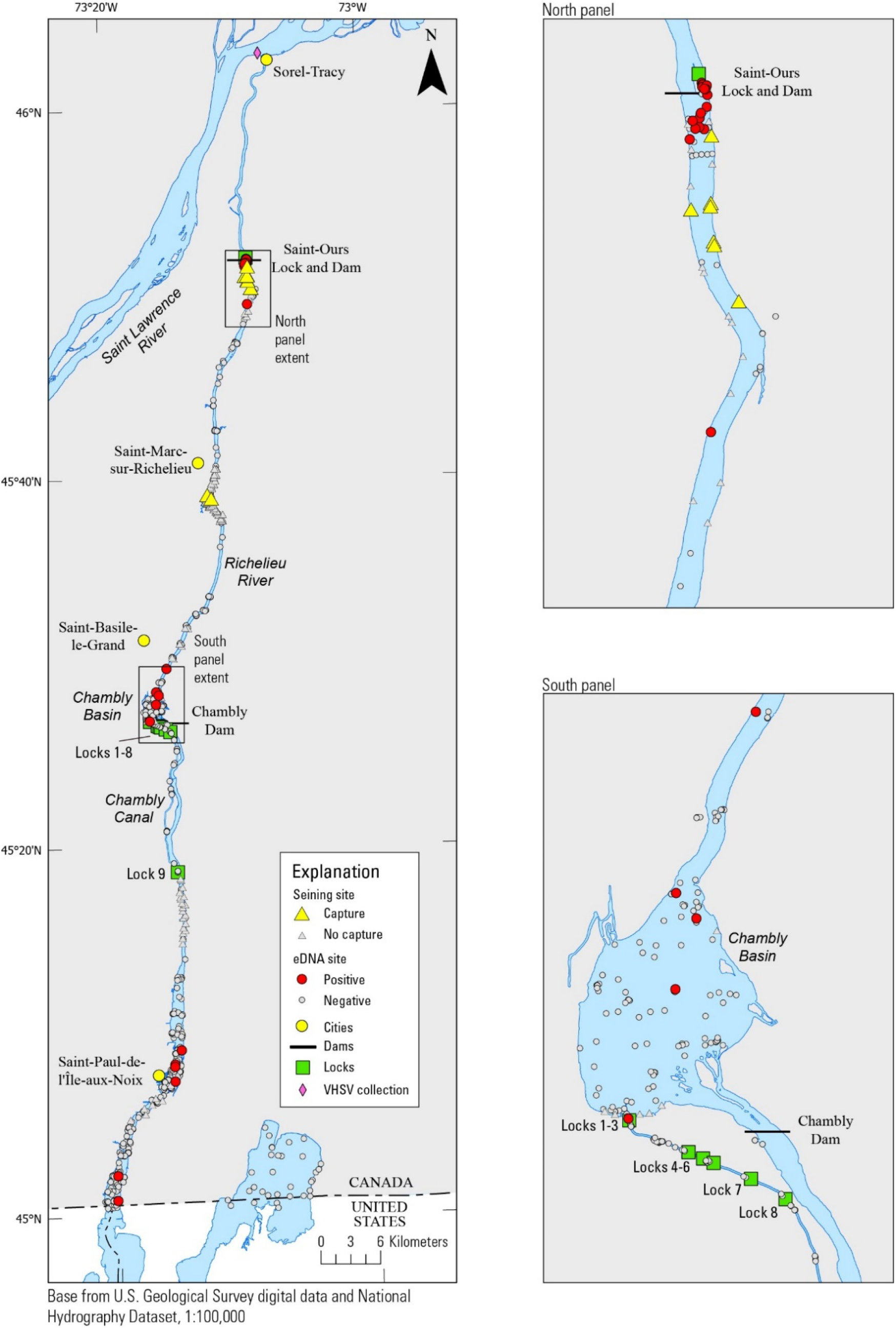
Environmental DNA (eDNA) and seining results from round goby (Neogobius melanostomus) surveys on the Richelieu River from 2021-2025.

In 2021, four of 70 eDNA samples produced positive detections of round goby DNA. Those four samples were located at Saint-Paul-de-l’île-aux-Noix (n=3) and near the Canadian side of the international border (n=1, 6 of 6 qPCR replicates positive) (Figure 5). In 2022, sampling was expanded to include the Saint-Ours area. Seven of 142 samples produced positive detections (from 1 to 6 qPCR replicates positive), all of which were located immediately upstream of the Saint-Ours Dam. In 2023, sampling was further expanded to include the Chambly Canal. Eleven of 176 samples produced positive detections (from 1 to 6 qPCR replicates positive), all located immediately upstream of the Saint-Ours Dam. In 2024, 3 of 141 samples produced positive detections (from 1 to 2 qPCR replicates positive). Those positives were located at Saint-Paul-de-l’île-aux-Noix (n=1), within the Chambly Basin (n=1), and downstream of the Chambly Basin (n=1). In 2025, 4 of 151 samples produced positive detections. Those positives were located at the Canadian side of the international border (n=1, 3 of 6 qPCR replicates positive) and in the Chambly Basin (n=3, 1 qPCR replicate positive for each sample).

Targeted beach seining for monitoring round goby distribution was conducted in 2023 and 2024 to investigate the positive eDNA samples from 2021 and 2022 in areas upstream of the known invasion front (Table 2). These efforts produced the first documentation of round goby upstream of the Saint-Ours Dam in 2023, with the capture of 3 individuals across 10 sites. In 2024, 43 individuals were captured across 12 sites in the same area, for a total of 46 round goby ranging from 18 to 51 mm in length. The targeted seining efforts in Chambly Basin, Saint-Basile-le-Grand, and Upper Richelieu produced no additional captures.

**Table 2.** Summary of beach seining efforts in the Richelieu River.

| Year | Location | River kilometers | Seining effort (number of sites) | Presence of round goby (number of sites) | Total number of individuals captured | Sampling activity |
| --- | --- | --- | --- | --- | --- | --- |
| 2022 | Saint-Marc-sur-Richelieu | 45-51 | 40 | 0 | 0 | Copper redhorse monitoring |
| 2023 | Saint-Marc-sur-Richelieu | 45-51 | 40 | 0 | 0 | Copper redhorse monitoring |
| 2023 | Upstream of Saint-Ours | 23-28 | 10 | 2 | 3 | Targeted round goby seining |
| 2023 | Upper Richelieu | 90-117 | 28 | 0 | 0 | Targeted round goby seining |
| 2024 | Saint-Marc-sur-Richelieu | 45-51 | 40 | 0 | 0 | Copper redhorse monitoring |
| 2024 | Upstream of Saint-Ours | 23-28 | 12 | 5 | 43 | Targeted round goby seining |
| 2024 | Saint-Basile-le-Grand | 63-66 | 6 | 0 | 0 | Targeted round goby seining |
| 2024 | Chambly Basin | 70-73 | 6 | 0 | 0 | Targeted round goby seining |
| 2025 | Saint-Marc-sur-Richelieu | 45-51 | 40 | 2 | 2 | Copper redhorse monitoring |

Standardized beach seining at 40 recurrent sites in the Saint-Marc-sur-Richelieu area as part of young-of-the-year copper redhorse monitoring produced no captures of round goby in 2022, 2023, or 2024. However, in 2025 a single round goby was captured at two different sites (36 mm and 42 mm in length). These captures make Saint-Marc-sur-Richelieu the southernmost location on the Richelieu River with physical captures of round goby.

During May 2024, 170 round goby from the confluence of the Richelieu River and Saint Lawrence River were tested for VHSV. The length of these individuals ranged from 25 to 94 mm. All specimens produced a result of non-detection for VHSV.

## Discussion

This study represents a substantial collaborative effort, involving multiple agencies across two countries and integrating two independent methods, traditional capture-based surveys and molecular detection using eDNA, for monitoring expansion of round goby populations in major pathways to Lake Champlain. We confirmed upstream movement of round goby in both the Champlain Canal and Richelieu River between 2021 and 2025.

However, the current distribution of round goby in the Champlain Canal and Richelieu River, and therefore the species’ true proximity to Lake Champlain, remains uncertain due to stark differences in the locations of eDNA detections and physical captures in each river system. In the Champlain Canal, physical capture data indicate that since being detected in the Hudson River in 2021 (Pendleton et al. 2022), round goby advanced upstream at least 5 km by June 2022 to the C1 dam. No physical captures occurred upstream of the C1 dam, but two eDNA samples collected in areas upstream of the C1 dam produced a positive detection – both in 2024 and immediately upstream of the C2 dam. Similarly, in the Richelieu River, the upstream-most physical capture of round goby occurred at Saint-Marc-sur-Richelieu (82 km from Lake Champlain) in 2025, while the upstream-most eDNA detections were near the Canadian side of the international border (4 km from Lake Champlain), occurring in 2021 and 2025. Additionally, eDNA sampling efforts within Lake Champlain proper using qPCR and metabarcoding (results not shown) during the same period produced no detections of round goby DNA.

Our results showing round goby eDNA detections occurring upstream of physical captures on both the Champlain Canal and Richelieu River are not surprising and likely arise from fundamental differences between the method types. First, physical capture methods and eDNA sampling do not seek the same target. Physical capture methods produce a result of presence or absence based on the attempted capture of a live animal, while eDNA samples are screened for a sequence of nucleotides determined to be unique to the target taxon. In both the Champlain Canal and Richelieu River, expansion towards Lake Champlain occurs in an upstream direction, so the upstream-most eDNA detections on each system cannot be easily attributed to downstream drift of DNA from any known round goby population. Nonetheless, numerous mechanisms that move DNA in aquatic ecosystems have been documented that could result in sample-level detection without site-level presence (Merkes et al. 2014; Darling et al. 2021; Stein et al. 2024; Xiong et al. 2024). Second, physical capture methods and eDNA methods represent vastly different spatial areas. Even when paired at the same time and location (as was frequently done in this study), physical capture methods such as electrofishing, trawling, and seining are spatially discrete sampling methods that exclusively represent only a small fraction of the available habitat, while eDNA samples integrate a larger and unknown area of habitat. Finally, although none of the eDNA control samples in this study produced evidence of field or laboratory-based contamination, the potential for false positives can never be completely ruled out with eDNA methods (Darling et al. 2021; Hutchins et al. 2022). Conversely, the physical capture methods may have low detection probabilities at low population densities and therefore are at elevated risk for producing false negative results (Wilcox et al. 2016). Considered alongside the entire body of research presented here, the upstream eDNA detections in both the Champlain Canal and the Richelieu River indicate that the species may be closer to Lake Champlain than indicated by capture data alone, and that additional targeted physical sampling is warranted to gain perspective on the methodological disagreement.

Our data do not decisively indicate whether round goby populations in the Champlain Canal and Richelieu River are carrying VHSV. In the Champlain Canal, one specimen met criteria for a positive detection, two specimens were classified as inconclusive, and 247 specimens were classified as non-detection. In the Richelieu River, all of the 170 tested specimens were classified non-detections. This finding is consistent with a 2011-2013 effort on parts of the Saint Lawrence River, bracketing the mouth of the Richelieu River, in which none of the 534 round goby or the 526 brown bullhead *Ameiurus nebulosus* tested positive for VHSV (MFFP 2016). In past outbreaks in New York, 27-100% of round goby have tested positive (Cornwell et al. 2012a; Getchell et al. 2019; Haws et al. 2025). However, the disease can be challenging to detect during dormant periods, when prevalence can be less than 5% (Cornwell et al. 2012b). The nearest confirmed VHSV detections to the southern end of Lake Champlain occurred in Cayuga Lake (Getchell et al. 2019) and Skaneateles Lake (Faisal et al. 2012), each more than 300 km from the Hudson River by water, while detections on the Saint Lawrence River have been confirmed approximately 223 km upstream from the mouth of the Richelieu River (Getchell et al. 2019). Our data, when interpreted within the context provided here, are insufficient for determining if the invasion front of round goby from the north or south is carrying VHSV towards Lake Champlain. Additional testing along the southern and northern invasion front would likely be necessary to decisively answer this question.

Our data indicate a strong influence of water temperature on habitat usage of round goby on the Champlain Canal. Round goby were rarely captured during near-shore electrofishing surveys when water temperature was below 10 °C. This observation likely indicates seasonal habitat use patterns rather than a change in the efficiency (capture probability) of the sampling method. Multiple studies have documented a seasonal movement pattern of round goby in which populations move into shallower habitats in the spring and retreat to deeper, more stable habitats in the fall. For example, in Cayuga Lake (Andres et al. 2020), Lake Michigan (Carlson et al. 2021), and Lake Ontario and the lower Niagara River (Pennuto et al. 2021), round goby almost exclusively occupied habitats less than 20 m during summer, but retreated to depths greater than 20-30 m in the fall as the water cooled. Similarly, seasonal migrations in and out of tributary streams to the Great Lakes have been observed (Pennuto et al. 2010; Blair et al. 2019; May et al. 2020).

Although most of these studies do not identify a water temperature associated with triggering seasonal migrations, Pennuto et al. (2021) identified 10 °C as the approximate water temperature at which spring and fall movements occurred. This aligns closely with our electrofishing data which indicate round goby on the Champlain Canal are more apt to occupy shallower and higher velocity habitats during the warm water period and partially but not completely retreat to deeper more stable habitats when water temperature drops below 10 °C. It is noteworthy, however, that the three most recent surveys at the Mohawk River at Peebles Island site produced relatively large catches at water temperatures ranging from 0.1-5.0 °C, indicating additional factors likely influence habitat usage.

Rates of upstream and downstream expansion of round goby populations into uncolonized habitats are highly sought-after information for natural resource managers. Our data provide some insight into the rate of upstream expansion in large rivers but also demonstrate the confounding role of barriers such as navigation infrastructure in producing these estimates. Expansion rates are estimated with the understanding that the exact dates of colonization are almost never known in any study area, but data generated from high-frequency, robust sampling programs are the most likely to detect expansion in a timely manner. In the Champlain Canal, capture data indicate that since being detected in the Hudson River in July 2021 (Pendleton et al. 2022), round goby advanced upstream from the confluence of the Mohawk and Hudson Rivers to the C1 dam by June 2022. This represents an upstream expansion rate of approximately 5 km in one year. However, between June 2022 and October 2025, physical capture data did not indicate any additional expansion, presumably due to the C1 dam preventing further movement. This is an interesting observation given the annual winter removal of the tainter gates at the C1 Dam, a practice that was initially identified by managing agencies as a potential pathway for further upstream expansion. The absence of clear evidence that this has occurred may indicate that concentrated flow through the gate bays during winter months may effectively form a velocity and/or thermal barrier to upstream passage of round goby. In the Richelieu River, round goby were first captured upstream of the Saint-Ours Dam in October 2023. At the conclusion of 2025, the upstream-most capture had occurred in the Saint-Marc-sur-Richelieu area, approximately 25 km upstream of the Saint-Ours Dam. However, the lack of regular physical sampling immediately upstream of Saint-Ours prior to 2023, and in the area downstream of Saint-Marc-sur-Richelieu during the study, prevents a precise estimate of expansion rate. Furthermore, the possibility of human-assisted movements of round goby in the Richelieu River (and elsewhere) must always be considered as possible alternatives to natural expansion, although the possession and use of live fish as bait has been completely prohibited in Quebec since 2017 (Canada 1990; MELCCFP 2026). Similarly, New York and Vermont each have specific baitfish regulations that prohibit use of round goby as fishing bait (NYSDEC 2022; Vermont General Assembly 2025). Together, the information from the Champlain Canal and Richelieu River cautiously indicate higher rates of upstream expansion than most other published estimates in which values typically ranged from 0.5 to 4 kilometers per year (Bronnenhuber et al. 2011; Kornis et al. 2012; Šlapanský et al. 2017). However, rates of upstream expansion from dammed and undammed systems likely cannot be compared, making it nearly impossible to estimate how quickly round goby will move upstream in heavily dammed systems like the Champlain Canal/upper Hudson River.

The data presented in this manuscript have already informed multiple management actions in the study area. These data are particularly relevant and time sensitive in the Champlain Canal where the network of locks and dams provide options for management action that might be impractical or impossible in less modified systems. As a result of knowledge gained from the monitoring efforts described here, a Trigger Action Response Plan (TARP) was developed as part of a round goby Rapid Response Plan for the Champlain Canal System (NYPA and NYSDEC 2023). The TARP specifies a series of adaptive management actions, including modifications to operation of the Champlain Canal, that escalate in intensity as round goby advance north towards Lake Champlain. For example, the finding that round goby catch rate in near-shore habitats was diminished below 10 °C was the basis for a decision to leave the tainter gates at the C1 dam in place for approximately one month longer than past precedent. The operations plan for this lock was modified in 2023 to stipulate that the gates would not be removed until water temperature fell below 4.4 °C (40 °F) for a minimum duration of 24 hours and would be reinstalled in the spring when the water temperature reached the same threshold (Canal Corporation 2023). Additionally, the two eDNA positives upstream of the C2 dam in 2024 caused immediate operational changes of locks C1-C4 (NYSDEC 2024) in accordance with the TARP. Lockages for recreational vessel traffic were restricted to a maximum of three per day in both northbound and southbound directions. Additionally, double flushing of locks prior to opening for boat traffic (already implemented at Locks C1 and C2) was extended to include Locks C3 and C4. Finally, the positive eDNA detections prompted NYSDEC to (a) initiate eDNA sampling at sites bracketing the positive location and (b) screen habitat type and access within each interlock pool between Locks C2 and C7 so that rapid sampling could be implemented in the future if additional positive detections occurred. Similarly, on the Richelieu River, the positive eDNA samples upstream of Saint-Ours in 2022 triggered targeted seining efforts in 2023, in which round goby were captured for the first time upstream of the dam. Together, these outcomes are significant because they are examples of early detection monitoring with eDNA (and physical collection) informing rapid responses in the form of management action. Moreover, our findings related to water temperature, expansion rates, and associated management actions, may have applications in other parts of North America and Europe facing current or imminent round goby invasion.

## Acknowledgements

The authors extend their appreciation to Barry Baldigo, Andrew Adams, Danny Skelton, Stevie Grace Wrighter, Zachary Kubsch, John Christopher, William Seamans, Lucas Reiter, and Zackary Althiser for field support, Aaron Maloy for eDNA analysis and interpretation, and Anna Haws for guidance on VHSV sampling and interpretation on the Champlain Canal. The authors also acknowledge the work and collaboration of M. Pimentel for helping with VSHV sampling in Quebec, Jade Larivière for the analysis of eDNA samples from the Chambly Canal, Ariane Thérien for assessing the specificity and sensitivity of the new round goby assay, and the MELCCFP field technicians that sampled on the Richelieu River: Yannicia Fréchette-Hudon, Mathew Labrèche Goudreau, Lucie Veilleux, Simon Bellefleur, Emmanuel Fortin, Rose Provençal-Lachance, Louis-Étienne Picard, Thierry Gariépy, Alexis Roy, Pierre-Alexis Drolet, and Timothé Therrien. This research was supported in part by funding from the Lake Champlain Basin Program/New England Interstate Water Pollution Control Commission and the New York State Department of Environmental Conservation. Any use of trade, firm, or product names is for descriptive purposes only and does not imply endorsement by the U.S. Government. The findings and conclusions in this article are those of the authors and do not necessarily represent the views of the U.S. Fish and Wildlife Service.

