## Supplemental Material for "Status of Round Goby Invasion Fronts in New York and Quebec: Implications for Lake Champlain"

^8^ Lake Champlain Basin Program/New England Interstate Water Pollution Control Commission

^9^ New York Power Authority, Environmental, Health & Safety

^10^ Cornell University, Department of Public and Ecosystem Health, College of Veterinary Medicine

^11^ New York State Department of Environmental Conservation, Bureau of Fisheries, Region 5

^12^ Fisheries and Oceans Canada, Maurice Lamontagne Institute

Any use of trade, firm, or product names is for descriptive purposes only and does not imply endorsement by the U.S. Government.

***Nmelanostomus_COI***

Primer development was done by Cecilia Hernandez from Bernatchez Lab at Université Laval, Québec, Canada.

Assay specificity and sensitivity was performed by Geneviève Parent and Ariane Thérien from Laboratory of Genomics, Demersal and Benthic Sciences Branch, Maurice Lamontagne Institute, Fisheries and Oceans Canada, Government of Canada

Table S1. Species-specific primers and probes for *Neogobius melanostomus*. Primer name indicates gene amplified, fragment length (bp).

| Scientific & Common name | Primer/Probe Gene | Sequence 5' > 3' | bp |
| --- | --- | --- | --- |
| *Neogobius melanostomus* | *COI_NEME_F* | CCCCTGGCAGGCAACTT | 65 |
| Round goby | *COI_NEME_R* | AGGTGAAGGGAGAAGATTGTCAAG |  |
|  | *COI_NEME_P* | CAGGAGCATCCGTCG |  |

1) Validation in silico

In silico validation was performed on all related species and species with the same distribution area as the targeted species. Table S2 provides the mitochondrial sequences of the cytochrome c oxidase subunit 1 (COI) gene from *Neogobius melanostomus* and non-target species that were used to assess the potential for non-specific amplification. A potential pairing of the primer on the non-target species was considered up to five mutations within the sequence.

**Table S2.** Information on mitochondrial sequences of the cytochrome c oxidase subunit 1 (COI) gene from *Neogobius melanostomus* and non-target species that were used to assess the potential for non-specific amplification. All accession numbers, or sequences, were downloaded from NCBI (2025). Not applicable (NA) is indicated when there are more than five mutations. Mutations are identified in bold and underlined.

| **Scientific name** | **Common name** | **Number of sequences** | **NCBI accession numbers** | **Origin** | **NEME_F (5’-3’)** | **NEME_R (5’-3’)** | **NEME_P (5’-3’)** |
| --- | --- | --- | --- | --- | --- | --- | --- |
| *Neogobius melanostomus* | Round goby | 84 | MW564535.1, MW564413.1, MW564376.1, MW564328.1, ON097309.1, ON097860.1, ON097852.1, ON097840.1, ON097798.1, ON097710.1, ON097668.1, ON097585.1, ON097545.1, ON097456.1, ON097427.1, ON097343.1, KX145236.1, KX145201.1, KX145135.1, KX145056.1, KX144993.1, KX586207.1, KX586206.1, HQ960511.1, KM373676.1, KM373670.1, KM373641.1, KM286769.1, KM286768.1, KM286767.1, KM286766.1, KM077846.1, KM077845.1, KM077844.1, KM077843.1, KM077842.1, KM077841.1, KM077840.1, KC501012.1, KC501011.1, KC501010.1, KC501009.1, KC501008.1, KC501007.1, KC501006.1, KC501005.1, KC501004.1, KC501003.1, KC501002.1, KC501001.1, KC501000.1, KC500999.1, KC500998.1, KC500997.1, KC500996.1, KC500995.1, KC500994.1, KC500993.1, KX247282.1, KX247281.1, KX247280.1, KX247279.1, KP247497.1, KP247496.1, KP247495.1, KY176544.1, MK439908.1, KR477229.1, KR477228.1, KR477073.1, KR477072.1, JQ623961.1, JX473750.1, JX473749.1, JX473748.1, JX473747.1, JX473746.1, JX473745.1, JX473744.1, JX473743.1, JX473742.1, JX473741.1, JX473740.1, EU524920.1, EU524919.1, EU524156.1, EU524155.1, EU524154.1, OM736846.1, OM736803.1, OM736802.1, OM736801.1, MW856909.1, MW856897.1, MW856835.1 | Black Sea, Georgia  Upper Austria, Austria  Ontario, Canada  South Moravia, Czechia  Bavaria, Germany  Lower Austria, Austria  Schleswig-Holstein, Germany  Kocaeli, Turkey  Tver Oblast, Russia  Yaroslavl Oblast, Russia  Moscow Oblast, Russia  Gulf of Gdańsk, Kaliningrad Oblast, Russia  Gulf of Gdańsk, Tricity, Poland  Istanbul, Turkey  Harju County, Estonia  Quebec, Canada  Wisconsin, United States  New York, United States | CCCCTGGCAGGCAACTT | AGGTGAAGGGAGAAGATTGTCAAG | CAGGAGCATCCGTCG |
| *Homo sapiens* | Human | 1 | NC012920.1 | England | CCC**T**T**A**GCAGG**G**AACT**A** | AGGTG**T**A**A**GGAGAAGATGG**T**TA**G**G | C**T**GGAGC**C**TCCGT**A**G |
| *Felis catus* | Domestic cat | 1 | NC001700.1 | ND | CCCCT**A**GC**C**GGCAAC**C**T | NA | CA**T**GAGC**T**TC**T**G**A**C**T** |
| *Canis familiaris* | Domestic dog | 1 | NC002008.4 | Korea | CC**A**CTGGC**T**GGCAA**TC**T | NA | CAGGAGCATCCGT**T**G |
| *Carassius auratus* | Goldfish | 5 | KF147851.1, KJ874428.1, KJ874430.1, KM659025.1, OK945938.1 | China  Oregon, United States | CC**T**CT**T**GCAGG**A**AAC**C**T | NA | CAGGAGCATC**A**GT**A**G |
| *Acipenser fulvescens* | Lake sturgeon | 10 | EU523878, EU524392, EU524393, EU524394, EU524395, EU524396, EU524397, KX145211, KX145284, KX145447 | Québec, Canada  Ontario, Canada | CC**G**CTGGC**G**GG**A**AAC**C**T | AGGTGAAGGGAGAA**A**AT**G**GT**T**A**G**G | C**G**GGAGC**C**TC**T**GT**A**G |
| *Acipenser oxyrinchus* | Atlantic sturgeon | 9 | EU523886, EU524398, EU524399, EU524400, EU524401, KX144976, KX145051, KX145066, KX145481 | Québec, Canada  Ontario, Canada | CC**A**CTGGC**G**GG**G**AACTT | NA | C**G**GGAGC**C**TC**T**GT**G**G |
| *Alosa aestivalis* | Blueback herring | 4 | KC015129, KX145025, KX145076, KX145570 | New-Brunswick, Canada | CCC**T**TGGCAGGCAA**T**CTT | NA | C**C**GGAGC**G**TCCGTCG |
| *Alosa pseudoharengus* | Alewife | 12 | CAISN096-12 (BOLD), EU523899, EU523900, EU524402, EU524403, PEC002-16 (BOLD), PEC004-16 (BOLD), PEC005-16 (BOLD), PEC007-16 (BOLD), PEC009-16 (BOLD), PEC010-16 (BOLD), PEC017-16 (BOLD) | Québec, Canada  Ontario, Canada | CCC**T**TGGCAGG**T**AA**T**CTT | AG**A**TGAAG**A**GAGAAGAT**A**GT**T**A**GA** | C**C**GGAGC**G**TCCGTCG |
| *Alosa sapidissima* | American shad | 10 | EU523901, EU523902, EU523903, EU524404, EU524405, EU524406, KX145092, KX145154, KX145379, KX145487 | Québec, Canada | CC**TT**TGGCAGGCAA**T**CTT | AG**A**TGAAG**A**GAGAAGAT**A**GT**T**A**G**G | C**C**GGAGCATCCGTCG |
| *Ambloplites rupestris* | Rock bass | 12 | EU523904, EU524407, EU524408, EU524409, EU524410, EU524411, EU524412, EU524413, EU524414, KX145339, KX145433, KX145606 | Québec, Canada  Ontario, Canada | CC**TT**T**A**GC**C**GGCAAC**C**T | AG**A**TGAAGGGAGAAGAT**G**GT**T**A**G**G | CAGG**T**GCATC**T**GTCG |
| *Ameiurus melas* | Black bullhead | 12 | EU523905, EU523906, EU524415, EU524416, EU524417, EU524418, EU524419, KX144985, KX145317, KX145369, KX145584, LTSM032-12 (BOLD) | Ontario, Canada | CC**A**CT**T**GC**T**GGCAAC**C**T | AGGTGAAG**T**GA**A**AAGAT**A**GT**T**A**GA** | CAGGAGC**C**TC**T**GT**A**G |
| *Ameiurus natalis* | Yellow bullhead | 12 | EU523908, EU524420, EU524421, EU524422, EU524423, EU524424, EU524425, KX145125, KX145200, KX145226, KX145424, KX145573 | Ontario, Canada | CC**A**CT**T**GC**T**GG**T**AAC**C**T | AGGTGAAG**T**GA**A**AAGAT**A**GT**T**AAG | C**G**GG**G**GC**C**TC**T**GT**A**G |
| *Ameiurus nebulosus* | Brown bullhead | 15 | EU523909, EU524426, EU524427, EU524428, EU524429, EU524430, EU524431, EU524432, EU524433, JX516987, JX517067, KX145148, KX145196, KX145343, KX145411 | Québec, Canada  Ontario, Canada | CC**A**CT**T**GC**T**GGCAAC**C**T | AGGTGAAG**T**GA**A**AAGAT**A**GT**T**A**GA** | CAGGAGC**C**TC**T**GT**A**G |
|  |  |  |  |  |  | AGGTGAAG**T**GA**A**AAGAT**A**GTCA**GA** |  |
| *Amia calva* | Bowfin | 8 | EU523910, EU524434, EU524435, KX145205, KX145217, KX145380, KX145432, KX145442 | Québec, Canada  Ontario, Canada | CC**T**CTGGCA**A**GCAAC**C**T | A**A**GTG**T**AGGGAGAAGAT**G**GT**T**AA**A** | CAGG**C**GCATC**A**GT**A**G |
| *Ammocrypta pellucida* | Eastern sand darter | 12 | EU523916, EU523915, EU523914, EU523913, EU523912, EU523911, KX145331, KX145591, KX145305, KX145150, KX145476, EU523917 | Québec, CA  Ontario, CA | CCCCTGGC**T**GGAAACTT | A**A**GTGAAG**T**GAGAAGATTGT**T**AA**A** | C**C**GGAGCATCCGT**T**G |
| *Anguilla rostrata* | American eel | 8 | EU523918, EU524440, EU524441, EU524442, KX145108, KX145210, KX145399, KX145504 | Québec, Canada  Ontario, Canada | CCCCTGGC**T**GG**A**AACTT | AGGTGAAG**T**GAGAA**A**ATTGTCA**G**G | C**C**GGAGCATC**T**GT**T**G |
| *Apeltes quadracus* | Fourspine stickleback | 10 | EU523919, EU524443, EU524444, EU524445, EU524446, KX144983, KX145268, KX145366, KX145387, KX145408 | Québec, Canada | CCCCTG**T**C**C**GG**A**AAC**C**T | NA | C**C**GG**C**GC**C**TC**A**GT**G**G |
| *Aplodinotus grunniens* | Freshwater drum | 9 | EU522441, EU522442, EU522443, EU522444, EU522445, KX145094, KX145163, KX145194, KX145212 | Québec, Canada  Ontario, Canada | NA | NA | CAGG**T**GC**T**TC**T**GT**T**G |
| *Campostoma anomalum* | Central stoneroller | 7 | KX145348.1, KX145324.1, KX145255.1, KX144997.1, KP013113.1, OM736851.1, NC_008102.1 | Ontario, Canada  Pennsylvania, United States | CC**A**CT**T**GC**G**GGCAA**TC**T | NA | C**G**GGAGCATC**A**GT**A**G |
| *Carpiodes cyprinus* | Quillback | 14 | EU524451, EU524452, EU524453, EU524454, EU524455, EU524456, EU524457, EU524458, EU524459, EU524460, EU524461, KX145067, KX145286, KX145578 | Québec, Canada  Ontario, Canada | C**A**CCTGGCAGG**AGT**CT**C** | AGGTG**G**AG**T**GAGAAGAT**A**GT**T**A**G**G | C**C**GGAGC**T**TC**T**GT**A**G |
| *Catostomus catostomus* | Longnose sucker | 26 | EU524462, EU524463, EU524464, EU524465, EU524466, EU524467, EU524468, EU524469, EU524470, EU524471, KR733317, KR733350, KR733351, KR733352, KR733353, KR733354, KR733355, KR733356, KR733357, KR733358, KR733359, KX145050, KX145102, KX145107, KX145430, KX145551 | Québec, Canada  Ontario, Canada | C**A**CCTGGCAGG**TGT**CT**C** | AGGTGAAG**A**GA**A**AAGAT**G**GT**T**AA**A** | C**C**GGAGC**T**TC**T**GT**A**G |
| *Catostomus commersonii* | White sucker | 20 | EU524476, EU524477, EU524478, EU524479, EU524480, EU524481, EU524482, EU524483, EU524484, KR733363, KR733364, KR733365, KR733378, KR733379, KX145116, KX145335, KX145540, KX145592, MNRFE003-14 (BOLD), MNRFE004-14 (BOLD) | Québec, Canada  Ontario, Canada | CC**A**CT**T**GC**G**GG**T**AA**T**CTT | AGGTGAAG**A**GA**A**AAGAT**G**GT**T**AAG | C**C**GGAGC**C**TC**T**GT**A**G |
| *Chrosomus eos* | Northern redbelly dace | 10 | EU525058, EU525059, EU525060, EU525061, EU525062, EU525063, KX145011, KX145248, KX145363, KX145388 | Québec, Canada  Ontario, Canada | NA | AGGTGAAGGGAGAA**A**ATTGT**T**A**G**G | CAGG**G**GCATC**A**GT**A**G |
|  |  |  |  |  |  | AGGTG**T**AGGGAGAAGAT**A**GTCA**G**G |  |
| *Chrosomus neogaeus* | Finescale Dace | 8 | EU525064, EU525065, EU525066, EU525067, EU525068, EU525069, EU525070, EU525071 | Québec, Canada  Ontario, Canada | CC**A**CT**T**GCAGG**T**AAC**C**T | AGGTGAAG**C**GAGAA**A**ATTGT**T**A**GA** | C**T**GGAGCATC**A**GT**A**G |
|  |  |  |  |  |  | AGGTGAAG**T**GAGAA**A**ATTGT**T**A**GA** | C**C**GGAGCATC**A**GT**A**G |
| *Coregonus artedi* | Cisco | 44 | EU523939.1, EU523940.1, EU523941.1, EU523942.1, EU523943.1, EU523944.1, EU523945.1, KP978018.1, KP978019.1, KP978020.1, KP978021.1, KP978022.1, KP978023.1, KP978024.1, KP978025.1, KP978026.1, KP978027.1, KP978028.1, KP978029.1, KP978030.1, KP978031.1, KR733386.1, KX145120.1, KX145160.1, KX145256.1, KX145323.1, KX145330.1, KX145353.1, KX145404.1, KX145505.1, KX145533.1, KX145581.1, KX145608.1, MT577062.1, MT577063.1, MT577064.1, MT577065.1, MT577066.1, MW856899.1, MW856900.1, OR814541.1, OR814542.1, OR814543.1, OR814544.1 | Ontario, Canada  Manitoba, Canada  New York, United States | CC**T**CTGGCAGGCAAC**C**T | NA | CAGG**G**GC**C**TCCGTCG |
| *Coregonus clupeaformis* | Lake whitefish | 62 | CYTC2364-12 (BOLD), CYTC2365-12 (BOLD), CYTC2366-12 (BOLD), CYTC2367-12 (BOLD), CYTC2368-12 (BOLD), CYTC2369-12 (BOLD), EU523957, EU523958, EU523959, JQ661482, JQ661483, JQ661484, JQ661485, JQ661486, JQ661487, KP978069, KP978070, KP978071, KP978072, KP978073, KP978074, KP978075, KP978076, KP978077, KP978078, KP978142, KR733387, KR733388, KR733389, KR733390, KR733391, KR733392, KR733393, KR733394, KR733395, KR733396, KR733397, KR733398, KR733399, KR733400, KR733401, KR733402, KR733403, KR733404, KR733405, KR733406, KR733407, KR733408, KR733409, KR733410, KR733411, KR733412, KR733413, KR733414, KR733415, KR733416, KR733417, KR733418, KR733419, KR733420, KR733421, KR733422 | Québec, Canada  Ontario, Canada | CC**T**CTGGCAGGCAAC**C**T | NA | CAGGAGC**C**TCCGTCG |
|  |  |  |  |  |  |  | CAGG**G**GC**C**TCCGTCG |
| *Cottus bairdii* | Mottled sculpin | 29 | EU522458, EU522457, EU522456, EU522455, EU522461, EU522460, EU522459, EU524506, EU524505, EU524504, EU524503, EU524502, EU524501, EU524500, EU524499, EU524498, EU524497, EU524496, EU524495, EU524494, EU524493, EU524492, EU524491, EU524490, KX145161, KX145329, KX145563, KX145193, EU523998 | Ontario, Canada  Quebec, Canada | CCCCTCGC**C**GG**A**AAC**C**T | AGGTGAAGGGAGAAGATTGT**T**A**G**G | CAGGAGC**C**TC**T**GT**T**G |
| *Cottus cognatus* | Slimy sculpin | 30 | EU524520, EU524519, EU524518, EU524517, EU524516, EU524515, EU524514, EU524513, EU524512, EU524511, EU524510, EU524509, EU524508, EU524507, MG421408, MG423581, MG422908, MG422298, MG422393, MG423337, MG422882, MT577118, MT577117, MT577116, MT577115, MT577114, OL766446, OL766447, OL769005, OL769275 | Québec, Canada  Ontario, Canada  New-Brunswick, Canada  Manitoba, Canada  Yukon, Canada,  Northwest Territories,  Canada | CCCCTTGC**C**GG**A**AAC**C**T | AGGTGAAGGGAGAAGATTGT**T**A**G**G | CAGGAGC**C**TC**T**GT**T**G |
| *Cottus ricei* | Spoonhead sculpin | 30 | MT577122, MT577119, MOBIL10235-19 (BOLD), MOBIL10315-19 (BOLD), MOBIL10768-20 (BOLD), JN025135, JN025136, EU524001, EU522463, EU522462, EU524521, KX145495, KX145393, KX145520, KX145517, MG421304, EF416989, MT577123, MT577121, MT577120, OL766645, MOBIL10200-19 (BOLD), MOBIL10210-19 (BOLD), MOBIL10219-19 (BOLD), MOBIL10312-19 (BOLD), MOBIL10313-19 (BOLD), MOBIL10314-19 (BOLD), MOBIL10316-19 (BOLD), MOBIL10383-19 (BOLD), MOBIL10744-20 (BOLD) | Alberta, Canada  Manitoba, Canada  Ontario, Canada  Quebec, Canada | CCCCTCGC**C**GG**A**AAC**C**T | AGGTGAAGGGAGAAGATTGTCA**G**G | CAGGAGC**C**TC**T**GT**T**G |
| *Couesius plumbeus* | Lake chub | 11 | EU524002, EU524523, EU524524, EU524525, EU524526, EU524527, EU524528, EU524529, EU524530, EU524531, KX145192 | Québec, Canada  Ontario, Canada | NA | AG**A**TGAAGGGAGAA**A**ATTGT**T**A**G**G | C**C**GG**C**GCATC**A**GT**A**G |
|  |  |  |  |  |  | AGGTGAAGGGAGAA**A**ATTGT**T**A**G**G |  |
| *Culaea inconstans* | Brook stickleback | 25 | BCFB057-06 (BOLD), BCFB060-06 (BOLD), EU524003, EU524532, EU524533, EU524534, EU524535, EU524536, EU524537, EU524538, JX516841, JX516917, JX516918, JX516975, JX517002, JX517014, JX517054, JX517082, JX517125, JX517129, JX517146, JX517164, KX145182, KX145565, KX145599 | Québec, Canada  Ontario, Canada | CC**T**CT**AT**C**T**GG**A**AACTT | AGGTGAAG**A**GA**A**AAGAT**G**GT**T**A**GA** | C**G**GGAGC**T**TC**A**GT**T**G |
|  |  |  |  |  | CCCCT**AT**C**C**GG**A**AACTT | AGGTGAAG**A**GA**A**AAGAT**G**GT**T**AA**A** | CAGGAGC**C**TC**A**GT**T**G |
|  |  |  |  |  | CCCCT**AT**C**T**GG**A**AACTT |  |  |
| *Cyprinella spiloptera* | Spotfin shiner | 16 | EU524548, EU524547, EU524546, EU524545, EU524544, EU524543, EU524542, EU524541, EU524540, EU524539, KX145239, KX145421, KX145326, KX144966, EU524004, EU524005 | Québec, Canada  Ontario, Canada | NA | NA | CAGGAGCATC**A**GT**A**G |
| *Cyprinus carpio* | Common carp | 11 | EU524549, EU524550, EU524552, EU524553, EU524554, EU524555, EU524556, KR733423, KX145028, KX145038, KX145139 | Québec, Canada  Ontario, Canada | CC**T**CT**T**GCAGG**G**AACTT | AGGTGAAG**T**GAGAA**A**ATTGT**T**A**G**G | CAGGAGCATC**A**GT**A**G |
| *Dorosoma cepedianum* | Gizzard shad | 9 | EU524558, EU524560, EU524561, EU524562, EU524563, EU524564, EU524565, EU524566, KX145014 | Ontario, Canada | NA | NA | C**C**GGAGCATCCGT**A**G |
| *Esox americanus americanus* | Redfin pickerel | 12 | EU524009, EU524569, EU524570, EU524571, EU524574, EU524575, EU524576, EU524577, JN025453, JN025454, JN025455, JX516898 | Québec, Canada | C**A**CCTGGCAGG**T**A**T**CT**C**  C**A**CCT**A**GCAGG**T**A**T**CT**C** | AGGTG**T**AG**A**GAGAAGATTGT**T**AAG  AGGTG**T**A**A**GGAGAAGATTGT**T**AAG | C**C**GG**C**GC**C**TCCGT**T**G |
| *Esox americanus vermiculatus* | Grass pickerel | 24 | EU524568, EU524572, EU524573, HQ556989, HQ556990, HQ557475, HQ557489, JN025462, JN025463, JN025464, JX516115, JX516514, JX516538, JX516662, JX516675, JX516808, JX516879, JX516965, JX516989, JX517042, JX517056, JX517070, JX517122, KX145297 | Ontario, Canada | C**A**CCTGGCAGG**T**A**T**CT**C**  C**A**CCT**A**GCAGG**T**A**T**CT**C** | AGGTG**T**AG**A**GAGAAGATTGT**T**AAG  AGGTG**T**A**A**GGAGAAGATTGT**T**AAG | C**C**GG**C**GC**C**TCCGT**T**G |
| *Esox lucius* | Northern pike | 21 | EU524010, EU524578, EU524579, EU524580, EU524581, EU524582, EU524583, EU524584, EU524585, EU524586, EU524587, EU524588, EU524589, EU524590, EU524591, EU524592, KX145022, KX145124, KX145189, KX145270, KX145489 | Québec, Canada  Ontario, Canada | CC**TT**TGGC**C**GG**A**AACTT | AGGTG**G**AG**A**GAGAA**A**AT**A**GT**T**AAG | CAGG**T**GC**T**TC**T**GT**A**G |
| *Esox masquinongy* | Muskellunge | 15 | EU524011, EU524593, EU524594, EU524595, EU524596, EU524597, EU524598, EU524599, EU524600, EU524601, EU524602, KX145156, KX145231, KX145548, KX145554 | Québec, Canada  Ontario, Canada | NA | NA | CAGG**G**GC**C**TC**T**GT**A**G |
| *Esox niger* | Chain pickerel | 14 | EU524012, EU524607, EU524608, EU524609, EU524610, EU524611, EU524612, HQ557067, HQ557068, HQ557069, KX145167, KX145199, KX145203, KX145480 | Québec, Canada | C**A**CCTGGCAGG**T**A**T**CT**C** | AGGTG**T**AG**A**GAGAAGATTGT**T**AAG | C**C**GG**C**GC**C**TCCGT**T**G |
| *Etheostoma blennioides* | Greenside darter | 12 | EU524019, EU524018, EU524017, EU524016, EU524015, EU524014, EU524013, KX145264, KX145173, KX145304, KX145157, KX144981 | Ontario, Canada | CCCCT**A**GC**T**GG**G**AAC**C**T | NA | CAGGAGCATCCGT**T**G |
| *Etheostoma exile* | Iowa darter | 16 | EU524029, EU524028, EU524027, EU524026, EU524025, EU524024, KX145392, KX145122, KX145071, EU524030, PPGB530-18 (BOLD), KP978063, KP978060, KP978064, KP978061, KP978059 | Québec, Canada  Ontario, Canada | CC**A**CTGGC**T**GG**A**AACTT | NA | C**C**GG**G**GCATC**T**GT**G**G |
| *Etheostoma caeruleum* | Rainbow darter | 11 | EU524023, EU524022, EU524021, EU524020, KX145225, KX145496, KX145450, KX145465, KX145315, MNRFE057-14 (BOLD), MNRFE074-14 (BOLD) | Ontario, Canada | CCCCT**A**GC**T**GG**G**AAC**C**T  CCGCT**A**GC**T**GG**T**AAC**C**T | NA | CAGGAGCATCCGT**T**G  C**C**GG**G**GCATCCGT**T**G |
|  |  |  |  |  |  | NA |  |
| *Etheostoma flabellare* | Fantail darter | 1  18 | EU524037, EU524036, EU524035, EU524034, EU524033, EU524032, EU524031, HQ557467, HQ557468, HQ557469, HQ557470, HQ557471, KX145383, KX144977, KX145349, KX145170, KX145204, EU524038 | Québec, Canada  Ontario, Canada | CC**A**CTGGC**T**GG**A**AACTT  CC**AT**TAG**CT**GG**T**AACTT | NA | C**C**GG**G**GCATC**T**GT**G**G  C**C**GG**G**GCATCCGT**T**G |
|  |  |  |  |  |  | NA |  |
| *Etheostoma microperca* | Least darter | 13 | EU524044, EU524043, EU524042, EU524041, EU524040, EU524039, KX145052, KX145222, KX145512, KX145254 | Ontario, Canada | CC**TT**TAGC**T**GG**A**AACTT | A**A**GTGAAGGGAGAA**A**ATTGT**A**A**G**G | C**G**GGAGCATC**T**GT**T**G |
| *Etheostoma nigrum* | Johnny darter | 142 | EU524046, EU524045, EU524056, EU524053, EU524052, EU524051, JX517023, JX517186, JX517166, JX516999, JX517168, JX517005, JX516913, JX516800, JX516842, JX516795, JX516792, JX516945, JX516957, JX516940, JX516901, JX517154, JX517027, JX516832, JX517055, JX516857, JX517084, JX516974, JX517163, JX517126, JX517073, JX517049, JX516833, JX517136, JX516818, JX516952, JX516740, JX516512, JX516328, JX516733, JX516315, JX516155, JX516179, JX516561, JX516764, JX516478, JX516341, JX516228, JX516264, JX516273, JX516376, JX516383, JX516221, JX516718, JX516286, JX516409, JX516240, JX516135, JX516475, JX516583, JX516783, JX516302, JX516177, JX516484, JX516168, JX516130, JX516646, JX516143, JX516247, JX516404, JX516156, JX516430, JX516151, JX516618, JX516559, JX516458, JX516267, JX516114, JX516415, JX516214, JX516747, JX516344, JX516686, JX516518, JX516650, JX516671, JX516294, JX516262, JX516705, JX516429, JX516630, JX516670, JX516300, JX516158, JX516655, JX516580, JX516118, JX516265, JX516313, JX516741, JX516162, JX516745, JX516734, JX516260, JX516340, JX516767, JX516513, JX516683, JX516490, JX516192, JX516205, JX516285, JX516771, JX516329, JX516653, JX516370, JX516688, JX516333, JX516178, JX516571, JX516311, JX516186, JX516137, JX516459, JX516456, JX516649, JX516261, JX516582, JX516275, JX516309, JX516615, JX516689, JX516331, JX516353, JX516510, JX516184, JX516427, JX516216, JX516212, JX516593, KX145406 | Québec, Canada  Ontario, Canada | CCCCT**A**GC**T**GG**G**AA**T**TT | NA | C**C**GG**G**GCATC**T**GT**T**G |
| *Etheostoma olmstedi* | Tessellated darter | 80 | EU524050, EU524049, EU524048, EU524047, EU524055, EU524054, JX516927, JX516794, JX517061, JX516978, JX516877, JX516959, JX516846, JX517085, JX517189, JX516897, JX517120, JX516888, JX517040, JX517124, JX517123, JX517170, JX516314, JX516604, JX516399, JX516317, JX516722, JX516661, JX516691, JX516291, JX516720, JX516529, JX516298, JX516113, JX516610, JX516434, JX516111, JX516460, JX516208, JX516282, JX516157, JX516570, JX516693, JX516743, JX516578, JX516245, JX516125, JX516723, JX516390, JX516739, JX516698, JX516433, JX516775, JX516780, JX516195, JX516778, JX516160, JX516292, JX516395, JX516358, JX516765, JX516131, JX516327, JX516190, JX516280, JX516526, JX516711, JX516284, JX516420, JX516180, JX516502, JX516440, JX516637, JX516283, JX516738, JX516631, JX516417, JX516337, JX516138, JX516715 | Québec, Canada  Ontario, Canada | CC**A**CT**A**GC**T**GG**G**AACTT | AGGTG**T**AGGGA**A**AAGAT**A**GT**T**AA**A** | C**C**GG**G**GCATCCGT**T**G |
| *Exoglossum maxillingua* | Cutlip minnow | 21 | NC_037015.1, OM736845.1, MT456071.1, MT455955.1, MT455703.1, KX145307.1, KX145262.1, KX145146.1, KX145119.1, KX145015.1, JN026614.1, JN026613.1, JN026612.1, JN026611.1, JN026610.1, JN026609.1, EU524616.1, EU524615.1, EU524614.1, EU524613.1, EU524057.1 | New York, United States  Pennsylvania, United States  Maryland, United States  Québec, Canada  Virginie, Canada | CC**A**CT**C**GCAGGCAA**TC**T | AGGTG**G**AG**A**GAGAAGATTGT**G**A**G**G | CAGG**G**GCATC**A**GT**A**G |
| *Fundulus diaphanus* | Banded killifish | 13 | EU524058, EU524620, EU524621, EU524622, EU524623, EU524624, EU524625, EU524626, KX145523, KX145547, KX145556, KX145559, KX145602 | Québec, Canada  Ontario, Canada | CC**A**CT**A**GCAGG**T**AA**T**TT | NA | CAGGAGC**T**TC**T**GT**A**G |
| *Fundulus notatus* | Blackstripe topminnow | 12 | EU524059, EU524060, EU524061, EU524062, EU524063, EU524064, EU524065, KX145017, KX145075, KX145105, KX145295, KX145460 | Ontario, Canada | CCCCT**A**GCAGG**A**AACTT | NA | CAGG**G**GC**T**TC**A**GT**A**G |
| *Gasterosteus aculeatus* | Threespine stickleback | 13 | EU524066, EU524631, EU524632, EU524633, EU524634, EU524635, EU524636, EU524637, EU524638, EU524639, KX145016, KX145412, KX145452 | Québec, Canada | CCCCT**CT**C**T**GG**G**AAC**C**T | AG**A**TGAAG**T**GA**A**AAGATTGT**T**A**G**G | CAGG**T**GC**T**TC**A**GTCG |
| *Gasterosteus wheatlandi* | Blackspotted stickleback | 4 | EU524067, EU524640, EU524641, EU524642 | Québec, Canada | NA | AG**A**TGAAG**A**GA**A**AAGATTGT**T**A**G**G | CAGG**T**GC**C**TC**A**GT**A**G |
|  |  |  |  |  |  | AG**A**TGAAG**A**GA**A**AAGATTGTCA**G**G |  |
| *Hiodon alosoides* | Goldeye | 4 | EU524648, EU524649, EU524650, EU524651 | Québec, Canada | CC**A**CT**A**GCAGG**T**AAC**C**T | NA | C**C**GG**C**GCATC**T**GT**T**G |
| *Hiodon tergisus* | Mooneye | 11 | EU524652, EU524653, EU524655, EU524656, EU524657, EU524658, EU524659, EU524660, EU524661, KX145397, KX145508 | Ontario, Canada  Québec, Canada | CC**G**CT**A**GCAGG**T**AAC**C**T | NA | C**C**GG**G**GCATCCGT**T**G |
| *Hybognathus argyritis* | Western silvery minnow | 12 | KX145413.1, KX145373.1, KX145243.1, KX145183.1, KX145012.1, EU524074.1, EU524073.1, EU524072.1, EU524071.1, EU524070.1, EU524069.1, EU522464.1 | Alberta, Canada  Missouri, États-Unis | CC**A**CT**T**GCAGG**T**AA**TC**T | AGGTG**G**AGGGAGAA**A**ATTGT**A**A**G**G | CAGGAGCATC**A**GT**A**G |
| *Hybognathus hankinsoni* | Brassy minnow | 17 | KX145202.1, KX145020.1, KX145013.1, JN026774.1, JN026773.1, JN026772.1, JN026771.1, JN026770.1, JN026769.1, JN026768.1, EU524081.1, EU524080.1, EU524079.1, EU524078.1, EU524077.1, EU524076.1, EU524075.1 | British Columbia, Canada  Québec, Canada  Michigan, United States  Nebraska, United States  Missouri, United States | CC**A**CTTGCAGGCAA**TC**T | A**A**GTG**G**AGGGAGAAGATTGT**A**A**G**G | CAGGGGCATC**A**GT**A**G |
| *Hybognathus placitus* | Plains minnow | 16 | JN026792.1, JN026791.1, JN026790.1, JN026789.1, EU524084.1, EU524083.1, EU524082.1, NC_056959.1, MW300341.1, MW300340.1, NC_056959.1 | Oklahoma, United States  Missouri, United States | CC**A**CTTGCAGG**T**AA**TC**T | AGGTGAAGGGAGAAGATTGT**A**A**G**G | CAGGAGCATC**A**GT**A**G |
| *Hybognathus regius* | Eastern silvery minnow | 19 | EU524085.1, EU524086.1, EU524662.1, EU524663.1, EU524664.1, EU524665.1, EU524666.1, GQ275151.1, KX145001.1, KX145043.1, KX145045.1, KX145405.1, KX145474.1, MG806595.1, MG806763.1, MT455025.1, MT455782.1, MT455797.1, MT456036.1 | Maryland, United States  Québec, Canada  South Carolina, United States | CC**A**CT**T**GCAGGCAA**TC**T | A**A**GTGAAGGGAGAAGATTGT**A**A**G**G | CAGG**G**GCATC**A**GT**A**G |
| *Ichthyomyzon castaneus* | Chestnut lamprey | 3 | EU524087, EU524088, EU524089 | Ontario, Canada | NA | AGGTG**T**AGGGAGAAGATTGT**T**AAG | CAGGAGC**C**TC**TA**T**T**G |
| *Ichthyomyzon fossor* | Northern brook lamprey | 6 | EU524090, EU524091, EU524093, EU524094, EU524095, EU524096 | Ontario, Canada | C**A**CCT**C**GC**T**GG**-**AA**T**TT | AGGTGAAGGGAGAAGATTGT**T**AAG | CAGGAGCATC**TA**T**T**G |
| *Ichthyomyzon unicuspis* | Silver lamprey | 9 | EU524097, EU524098, EU524099, EU524100, EU524101, EU524102, EU524103, EU524104, EU524105 | Ontario, Canada  Québec, Canada | C**A**CCT**C**GC**T**GG**-**AA**T**TT | AGGTGAAGGGAGAAGATTGT**T**AAG | CAGGAGCATC**TA**T**T**G |
| *Ictalurus punctatus* | Channel catfish | 28 | EU524106, EU524676, EU524677, EU524678, EU524679, EU524680, EU524681, EU524682, EU524683, EU524684, EU524685, EU524686, KX145159, KX145299, KX145334, KX145376, KX145470, LTSM015-12 (BOLD), LTSM019-12 (BOLD), LTSM150-12 (BOLD), LTSM159-12 (BOLD), LTSM196-12 (BOLD), LTSM215-12 (BOLD), LTSM217-12 (BOLD), LTSM219-12 (BOLD), LTSM222-12 (BOLD), LTSM329-12 (BOLD), LTSM376-12 (BOLD) | Québec, Canada  Ontario, Canada | CC**T**CT**T**GC**C**GGCAAC**C**T | AG**A**TGAAGGGA**A**AAGAT**A**GT**T**AA**A** | CAGG**G**GC**C**TCCGT**A**G |
|  |  |  |  |  |  |  | CAGGAGC**C**TCCGT**A**G |
| *Labidesthes sicculus* | Brook silverside | 25 | EU524108.1, EU524689.1, EU524690.1, EU524691.1, EU524692.1, EU524693.1, EU524694.1, EU524695.1, EU524696.1, EU524697.1, EU524698.1, JN026925.1, JN026926.1, JN026927.1, JN026928.1, JN026929.1, JN026930.1, KF930009.1, KX145096.1, KX145145.1, KX145367.1, KX145457.1, KX145546.1, MW856879.1, MW856908.1 | Québec, Canada  Ontario, Canada  New York, United States  Wisconsin, United States  Illinois, United States  Kansas, United States  South Carolina, United States | CC**T**CT**AT**CAGGCAA**TC**T | A**A**GTGAAGGGAGAAGAT**A**GT**T**AA**A** | C**C**GG**G**GCATCCGT**A**G |
|  |  |  |  |  | C**A**CCT**A**GCAGGCA**TT**T**C** | AGGTG**G**AGGGAGAA**A**AT**G**GT**T**AA**A** | C**C**GGAGCATCCGT**A**G |
|  |  |  |  |  | **TT**CCT**A**GCAGGC**TT**CTT |  | CAG**C**GAGCATCCGTC**T** |
| *Lepisosteus oculatus* | Spotted gar | 4 | EU524699, KX145147, KX145384, KX145427 | Ontario, Canada | CCCCT**A**GC**CA**GCAAC**C**T | NA | CAGGAGCATC**A**GT**T**G |
| *Lepisosteus osseus* | Longnose gar | 7 | EU524119, EU524120, EU524121, KX144975, KX145082, KX145372, KX145443 | Québec, Canada  Ontario, Canada | CCCCTGGC**TA**GCAA**TC**T | A**A**GTGAAGGGAGAA**A**AT**G**GT**T**A**GA** | CAGGAGCATC**A**GT**T**G |
| *Lepomis cyanellus* | Green sunfish | 13 | EU524705, EU524706, EU524707, EU524708, EU524709, EU524710, EU524711, EU524712, EU524713, KX145607, OP418481, OP418482, OP418483 | Ontario, Canada | CC**T**CT**C**GC**G**GGCAA**TC**T | NA | CAGG**G**GCATCCGT**G**G |
| *Lepomis gibbosus* | Pumpkinseed | 19 | EU524123, EU524714, EU524715, EU524716, EU524717, EU524718, EU524719, EU524720, EU524721, EU524722, EU524723, EU524724, EU524725, KX145030, KX145090, KX145175, KX145233, KX145351, KX145396 | Québec, Canada  Ontario, Canada | CC**T**CT**C**GC**C**GGCAAC**C**T | NA | C**C**GGAGCATCCGT**T**G |
| *Lepomis macrochirus* | Bluegill | 10 | EU524732, EU524733, EU524734, EU524735, EU524736, EU524737, EU524738, EU524739, EU524740, EU524741 | Ontario, Canada | CC**T**CT**C**GC**T**GG**T**AAC**C**T | AG**A**TG**C**AGGGAGAAGAT**A**GT**A**A**G**G | CAGGAGCATC**A**GTCG |
| *Lepomis megalotis* | Longear sunfish | 10 | EU524124, EU524742, EU524743, EU524744, EU524745, KX145130, KX145190, KX145364, KX145560, KX145582 | Québec, Canada  Ontario, Canada | CC**T**CT**C**GC**C**GGCAAC**C**T | NA | CAGG**G**GC**C**TCCGTCG |
| *Lota lota* | Burbot | 10 | EU524749, EU524750, EU524751, EU524752, EU524753, EU524754, EU524755, EU524756, EU524757, KX145483 | Québec, Canada  Ontario, Canada | CC**T**CT**A**GCAGGCAA**T**CTT | A**AA**TG**C**AGGGAGAAGAT**A**GT**A**A**G**G | C**T**GG**G**GC**T**TC**T**GT**T**G |
| *Lythrurus umbratilis* | Redfin shiner | 1 | NC_033935.1 | ND | CC**A**CT**TT**CAGG**T**AAC**C**T | AGGTG**G**AGGGAGAA**A**ATTGT**G**A**G**G | CAGGAGC**G**TC**A**GT**A**G |
| *Luxilus cornutus* | Common shiner | 42 | EU524126, EU524127, EU524765, EU524766, EU524767, EU524768, EU524769, EU524770, EU524771, EU524772, EU524773, EU524774, EU524775, EU524776, EU524777, EU524778, EU524779, EU524780, EU524781, EU524782, EU524783, EU524784, EU524785, EU524786, KX145171, KX145281, KX145437, KX145439, KX145445, MG423463, OP418466, OP418467, OP418468, PV369172.1, PV369173.1, PV369174.1, PV369182.1, PV369185.1, PV624624.1, PV624625.1, PV624626.1, SWBRP669-18 (BOLD) | Ontario, Canada  Québec, Canada | CC**G**CT**C**GCAGG**T**AAC**C**T | AG**A**TG**T**AGGGAGAA**A**ATTGT**A**A**G**G | CAGGAGCATC**A**GT**A**G |
| *Macrhybopsis storeriana* | Silver chub | 1 | NC_030485.1 | ND | CC**A**CT**T**GCAGG**T**AAC**C**T | AGGTGAAG**A**GA**A**AAGATTGT**G**A**G**G | CAG**C**GAGCATCC**A**TC**T** |
| *Margariscus margarita* | Allegheny pearl dace | 10 | EU524803, EU524804, EU524805, EU524806, EU524807, EU524808, EU524809, KX145033, KX145100, KX145600 | Québec, Canada  Ontario, Canada | CC**A**CT**C**GCAGG**T**AA**TC**T | A**A**GTGAAGGGA**A**AAGATTGT**T**A**G**G | C**C**GG**G**GCATC**A**GT**T**G |
| *Micropterus dolomieu* | Smallmouth bass | 23 | EU524131, EU524810, EU524811, EU524812, EU524813, EU524815, EU524816, EU524817, EU524818, EU524819, EU524820, EU524821, EU524822, EU524823, EU524824, EU524825, EU524826, EU524827, EU524828, KX145308, KX145375, KX145562, KX145588 | Québec, Canada  Ontario, Canada | CC**T**CT**T**GC**C**GGCAAC**C**T | AG**A**TGAAG**A**GAGAAGAT**G**GT**T**A**G**G | CAGGAGCATCCGT**T**G |
| *Micropterus salmoides* | Largemouth bass | 18 | EU524132, EU524829, EU524830, EU524831, EU524832, EU524833, EU524834, EU524835, EU524836, EU524837, EU524838, KX145023, KX145078, KX145132, KX145274, KX145587, LTSM165-12 (BOLD), MG423016 | Québec, Canada  Ontario, Canada | CC**T**CT**T**GC**C**GGCAAC**C**T | AGGTGAAG**A**GAGAAGAT**G**GT**T**A**G**G | CAGGAGCATCCGT**T**G |
| *Morone americana* | White perch | 9 | EU524133, EU524134, EU524135, EU524136, EU524137, EU524138, EU524139, KX145101, KX145215 | Québec, Canada  Ontario, Canada | CC**A**CT**T**GCA**A**G**T**AAC**C**T | AG**A**TG**G**AGGGAGAA**A**ATTGT**T**AA**A** | CAGGAGCATCCGT**A**G |
| Morone chrysops | White bass | 9 | EU524140, EU524141, EU524142, KX144971, KX145191, KX145448, LTSM040-12 (BOLD), LTSM131-12 (BOLD), LTSM132-12 (BOLD) | Ontario, Canada | CCCCT**T**GCA**A**GCAAC**C**T | AGGTG**G**AG**A**GA**A**AA**A**ATTGT**T**AAG | CAGG**T**GCATC**T**GT**A**G |
| *Morone saxatilis* | Striped bass | 3 | EU524143, EU524144, EU524145 | Québec, Canada | CCCCT**T**GCA**A**GCAAC**C**T | AGGTG**G**AG**A**GA**A**AA**A**ATTGT**T**AAG | CAGG**T**GCATC**T**GT**A**G |
|  |  |  |  |  | CC**T**CT**T**GCA**A**GCAAC**C**T | AGGTG**G**AGGGAGAA**A**ATTGT**T**A**G**G | CAGG**T**GCATC**T**GT**A**G |
| *Moxostoma anisurum* | Silver redhorse | 15 | EU524146, EU524846, EU524847, EU524848, EU524849, EU524850, EU524851, EU524852, EU524853, EU524854, EU524855, KX145247, KX145258, KX145435, KX145486 | Québec, Canada  Ontario, Canada | CCCCT**C**GC**G**GGCAA**T**CTT | NA | C**C**GGAGC**C**TC**T**GT**A**G |
| *Moxostoma carinatum* | River redhorse | 9 | EU524147, EU524148, EU524856, EU524857, EU524858, EU524859, EU524860, KX145093, KX145198 | Québec, Canada  Ontario, Canada | CCCCT**C**GC**G**GGCAA**T**CTT | NA | C**C**GGAGC**C**TC**T**GT**A**G |
| *Moxostoma duquesnei* | Black redhorse | 11 | EU524861, EU524862, EU524863, EU524864, EU524865, EU524866, KX145006, KX145087, KX145455, KX145488, KX145492 | Ontario, Canada | CCCCT**T**GC**T**GGCAA**T**CTT | NA | C**T**GGAGC**C**TC**T**GT**A**G |
| *Moxostoma erythrurum* | Golden redhorse | 13 | EU524867, EU524868, EU524869, EU524870, EU524871, EU524872, EU524873, EU524874, EU524875, EU524876, KX145327, KX145409, KX145507 | Ontario, Canada | CCCCT**C**GC**G**GGCAA**T**CTT | NA | C**C**GGAGC**C**TC**T**GT**A**G |
| *Moxostoma hubbsi* | Copper redhorse | 12 | EU524877, EU524878, EU524879, EU524880, EU524881, EU524882, EU524883, EU524884, EU524885, EU524886, EU524887, EU524888 | Québec, Canada | CCCCT**C**GC**G**GGCAA**T**CTT | NA | C**C**GGAGC**C**TC**T**GT**A**G |
| *Moxostoma macrolepidotum* | Shorthead redhorse | 20 | EU524149, EU524889, EU524890, EU524891, EU524892, EU524893, EU524894, EU524895, EU524896, EU524897, EU524898, EU524899, EU524900, EU524901, EU524902, EU524903, KX145069, KX145338, KX145484, KX145493 | Québec, Canada  Ontario, Canada | CCCCT**C**GC**G**GGCAA**T**CTT | NA | C**C**GGAGC**C**TCCGT**A**G |
| Moxostoma valenciennesi | Greater redhorse | 14 | EU524150, EU524904, EU524905, EU524906, EU524907, EU524908, EU524909, EU524910, EU524911, EU524912, KX145359, KX145497, KX145532, KX145564 | Québec, Canada  Ontario, Canada | CCCCT**C**GC**G**GGCAA**T**CTT | NA | C**C**GGAGC**C**TC**T**GT**A**G |
| *Myoxocephalus thompsonii* | Deepwater sculpin | 9 | EU524918, EU524917, EU524916, EU524915, EU524914, KX145213, KX145111, KX145164, KX145285, | Manitoba, Canada  Ontario, Canada | CCCCT**T**GC**C**GG**A**AAC**C**T | AG**A**TG**T**A**A**GGAGAAGATTGT**T**A**G**G | C**G**GGAGC**C**TC**T**GT**T**G |
| *Myoxocephalus quadricornis* | Fourhorn sculpin | 20 | EU524913, MG422966, MG421040, MG421302, MG423121, MG421539, MG421145, MG421255, MG422195, MG422697, MG421164, DSFIB010-06 (BOLD), MG422128, MG421777, MG421699, MG421387, MG421069, MG421042, MG421784, WHBI472-23 (BOLD), | Ontario, Canada,  Manitoba, Canada  Nunavut, Canada | CCCCT**T**GC**C**GG**A**AAC**C**T | AG**A**TG**T**A**A**GGAGAAGATTGT**T**A**G**G | C**G**GGAGC**C**TC**T**GT**T**G |
| *Nocomis biguttatus* | Hornyhead chub | 7 | EU524157.1, EU524158.1, EU524159.1, EU524160.1, EU524921.1, EU524922.1, EU524923.1 | Manitoba, Canada  Ontario, Canada | CC**T**CT**T**GCAGG**T**AAC**C**T | NA | CAGGAGCATC**G**GT**A**G |
| *Nocomis micropogon* | River chub | 2 | NC_042391.1, MW856863.1 | New York, United States  Pennsylvania, United States | CC**T**CT**C**GCAGG**T**AAC**C**T | NA | CAGGAGCATC**A**GT**A**G |
| *Notemigonus crysoleucas* | Golden shiner | 14 | EU524933, EU524934, EU524935, EU524936, EU524937, EU524938, EU524939, EU524940, JX516121, JX516207, JX516308, JX516363, JX516375, JX516450 | Québec, Canada | CCCCT**C**GCAGG**T**AA**TC**T | AGGTG**G**AG**T**GAGAAGAT**C**GT**T**AAG | CAGG**C**GC**G**TC**A**GT**A**G |
|  |  |  |  |  |  | AGGTG**G**AG**T**GAGAAGATTGT**T**AAG | CAGG**C**GCATC**A**GT**A**G |
|  |  |  |  |  |  |  | CAGGAGCATC**A**GT**A**G |
| *Notropis photogenis* | Silver shiner | 20 | KX145473.1, KX145306.1, KX145278.1, KX145138.1, KX145035.1, JN027633.1, JN027631.1, JN027632.1, JN027630.1, JN027629.1, AY116187.1, EU525015.1, EU525014.1, EU525013.1, EU525012.1, EU525011.1, EU525010.1, EU525009.1, EU525008.1, EU525007.1 | Ontario, Canada  West Virginia, United States  Pennsylvania, United States  Ohio, United States | NA | AGGTG**T**AG**A**GAGAA**A**ATTGT**G**A**G**G  AGGTGTAG**A**GAGAA**A**ATTGT**G**A**G**G | CAGGAGCATC**A**GT**A**G |
| *Notropis volucellus* | Mimic shiner | 3 | NC_080906.1, MW856882.1, OR552078.1 | Missouri, United States  Minnesota, États-Unis | CC**A**CT**T**GC**G**GG**T**AAC**C**T | AGGTGAAG**A**GAGAA**A**ATTGT**T**A**G**G | CAGGAGC**G**TC**A**GT**A**G |
| *Notropis texanus* | Weed shiner | 15 | KX145534.1, KX145511.1, KX145394.1, JN027728.1, JN027727.1, JN027726.1, JN027725.1, JN027724.1, JN027723.1, JN027722.1, JN027721.1, JN027720.1, JN027719.1, HQ937028.1, EU524182.1, s | Alabama, United States  Floride, United States  Georgia, United States  Manitoba, Canada | CC**A**CT**T**GCAGG**T**AAC**C**T | AGGTG**G**AGGGAGAAGATTGT**A**A**G**G | CAGG**G**GCATC**A**GT**A**G |
| *Notropis stramineus* | Sand shiner | 16 | EU524179.1, EU524180.1, EU524181.1, EU525024.1, EU525027.1, EU525030.1, HM179632.1, HM179633.1, HM179635.1, HM179637.1, JN027579.1, JN027581.1, JN027708.1, KF930192.1, KX145114.1, KX145522.1 | Québec, Canada  Indiana, United States  Pennsylvania, United States  Ontario, Canada  South Dakota, United States  West Virginia, United States | CC**A**CT**T**GCAGGCAA**TC**T | AGGTG**G**AG**A**GA**A**AAGATTGT**G**A**G**G | CAGG**G**GCATC**T**GT**A**G |
| *Notropis rubellus* | Rosyface shiner | 13 | EU524178.1, EU525016.1, EU525017.1, EU525018.1, EU525019.1, EU525020.1, EU525021.1, EU525022.1, KX145061.1, KX145350.1, KX145423.1, KX145482.1, KX145579.1 | Ontario, Canada | CC**A**CT**TT**CAGG**A**AAC**C**T | AGGTG**G**AG**A**GAGAAGATTGT**T**A**G**G | CAGGAGCATC**A**GT**A**G |
| *Notropis procne* | Swallowtail shiner | 15 | MT456222.1, MT455763.1, MT456156.1, MT455749.1, MT455324.1, MT455239.1, MT455172.1, MT455041.1, JN027639.1, JN027638.1, JN027637.1, JN027636.1, JN027635.1, HQ557391.1, HQ557390.1 | Maryland, United States  South Carolina, United States  Virginia, United States | CC**A**CT**T**GCAGG**T**AA**TC**T | AGGTGAAG**A**GAA**A**AGATTGT**G**A**G**G | CAGGAGCATCCGT**A**G |
| *Notropis hudsonius* | Spottail shiner | 7 | MT577131.1, MT577132.1, MT577133.1, MT577134.1, NC_037014.1, OM736809.1, OM736832.1 | Lake Winnipeg, Canada  Lale Michigan, United States  Lake Superior, United States | CC**A**CT**T**GCAGGCAA**TC**T | AGGTGAAG**A**GAGAA**A**ATTGT**G**A**G**G | C**G**GG**C**GCATC**A**GT**A**G |
| *Notropis heterodon* | Blackchin shiner | 16 | KX145586.1, KX145026.1, KX145019.1, KF930188.1, EU524991.1, EU524990.1, EU524989.1, EU524988.1, EU524987.1, EU524986.1, EU524985.1, EU524984.1, EU524983.1, EU524982.1, EU524981.1, EU524175.1 | Michigan, United States  Ontario, Canada  Québec, Canada | CC**A**CT**T**GCAGG**T**AA**TC**T | AGGTG**A**AG**A**GAAAAGATTGT**A**A**G**G | CAGGAGCATC**A**GT**A**G |
| *Notropis heterolepis* | Blacknose shiner | 24 | EU524992.1, EU524993.1, EU524994.1, EU524995.1, EU524996.1, EU524997.1, EU524998.1, EU524999.1, HQ557114.1, JN027547.1, JN027548.1, KX145110.1, KX145172.1, KX145230.1, KX145310.1, KX145346.1, KX145401.1, KX145603.1, MG570414.1, NC_037009.1, OP418458.1, OP418459.1, OP418460.1, OP418461.1 | Illinois, United States  New-Brunswick, Canada  Ontario, Canada  Québec, Canada | CC**A**CT**T**GCAGG**T**AAC**C**T | AGGTG**G**AG**A**GAGAAGATTGT**G**A**G**G | CAGGAGCATC**A**GT**A**G |
| *Notropis buchanani* | Ghost shiner | 24 | KX145342.1, KX145097.1, KX145004.1, KX144978.1, JN027486.1, JN027485.1, JN027484.1, JN027483.1, JN027482.1, JN027481.1, JN027480.1, JN027479.1, KP978065.1, EU524980.1, EU524979.1, EU524978.1, EU524977.1, EU524976.1, EU524975.1, EU524974.1, EU524973.1, EU524972.1, EU524971.1, EU524970.1 | Alabama, United States  Illinois, United States  Ontario, Canada  Texas, United States | CC**A**CT**T**GC**G**GG**T**AAC**C**T | AGGTGAAG**A**GAGAA**A**ATTGT**T**A**G**G | CAGGAGC**G**TC**A**GT**A**G |
| *Notropis boops* | Bigeye shiner | 13 | HQ579029.1, JN027454.1, JN027455.1, JN027456.1, JN027457.1, JN027458.1, JN027459.1, JN027460.1, JN027461.1, JN027462.1, JN027463.1, KF930187.1, MK037233.1 | Arkansas, United States  Oklahoma, United States | CC**A**CT**T**GC**G**GGCAA**TC**T | AGGTG**G**AG**A**GAGAAGATTGT**A**A**G**G | C**G**GG**G**GCATC**A**GT**A**G |
| *Notropis blennius* | River shiner | 11 | KX145461.1, KX145232.1, KX145064.1, KX145057.1, KX144989.1, JN027453.1, JN027452.1, JN027451.1, JN027450.1, JN027449.1, HQ557210.1 | Arkansas, United States  Illinois, United States  Manitoba, Canada | CC**A**CT**C**GCAGGCAAC**C**T | AGGTG**G**AGGGAGAAGATTGT**A**A**G**G | CAGG**C**GCATC**A**GT**A**G |
| *Notropis bifrenatus* | Bridle shiner | 17 | EU524172.1, EU524173.1, EU524174.1, EU524963.1, EU524964.1, EU524965.1, EU524966.1, EU524967.1, EU524968.1, EU524969.1, JN027448.1, KF930186.1, KX145301.1, KX145438.1, KX145459.1, KX145535.1, KX145593.1 | New Hampshire, United States  New Jersey, United States  Ontario, Canada,  Quebec, Canada | CC**A**CT**T**GCAGG**T**AAC**C**T | NA | CAGGAGCATC**A**GT**A**G |
| *Notropis anogenus* | Pugnose shiner | 20 | EU524162.1, EU524163.1, EU524164.1, EU524165.1, EU524166.1, EU524167.1, EU524168.1, EU524941.1, EU524942.1, EU524943.1, EU524944.1, EU524945.1, EU524946.1, EU524947.1, EU524948.1, EU524949.1, KX145178.1, KX145187.1, KX145296.1, KX145298.1 | Ontario, Canada | NA | AGGTGAAG**A**GAGAAGATTGT**A**A**G**G | CAGGAGCATC**A**GT**A**G |
| *Notropis atherinoides* | Emerald shiner | 7 | MT577130.1, MT577129.1, MT577128.1, MT577127.1, MW856878.1, MG570455.1, MG570456.1 | Manitoba, Canada  Wisconsin, United States  New York, United States | CC**A**CT**T**TCAGG**A**AAC**C**T | AGGTG**G**AG**A**GAGAAGATTGT**T**A**G**G | CAGGAGC**G**TC**A**GT**A**G |
| *Noturus flavus* | Stonecat | 5 | EU525039, EU525040, EU525041, EU525042, JN027792 | Québec, Canada | CCCCT**T**GC**C**GG**A**AAC**C**T | AGGTG**G**AGGGAGAAGAT**G**GT**T**AA**A** | CAGGAGC**C**TC**T**GT**A**G |
| *Noturus gyrinus* | Tadpole madtom | 10 | EU525044, EU525045, EU525046, EU525047, EU525048, EU525049, EU525050, EU525051, EU525052, KX145282 | Québec, Canada  Ontario, Canada | CCCCT**T**GC**C**GG**A**AAC**C**T | AGGTG**G**AGGGAGAAGAT**G**GT**T**AA**A** | CAGGAGC**C**TC**T**GT**A**G |
| *Noturus insignis* | Margined madtom | 4 | EU524186, EU524187, EU524188, EU524189 | Ontario, Canada | CCCCT**T**GC**C**GG**A**AAC**C**T | AGGTG**G**AGGGAGAAGAT**G**GT**T**AA**A** | CAGG**G**GC**C**TC**T**GT**A**G |
| *Noturus miurus* | Brindled madtom | 6 | EU525053, KX145010, KX145029, KX145356, KX145566, KX145571 | Ontario, Canada | CCCCT**T**GC**C**GG**A**AA**T**CT | AGGTGAAGGGAGAAGAT**G**GT**T**A**GA** | CAGGAGC**C**TC**T**GT**A**G |
| *Noturus stigmosus* | Northern madtom | 2 | EU525054, EU525055 | Ontario, Canada | CCCCT**C**GC**C**GG**A**AAC**C**T | AGGTGAAGGGAGAAGAT**G**GT**T**A**GA** | C**T**GGAGC**C**TC**T**GT**A**G |
| *Oncorhynchus kisutch* | Coho salmon | 2 | KX144987, KX145490 | Ontario, Canada | CC**T**CTGGC**C**GGCAAC**C**T | A**AA**TGAAGGGAGAAGAT**A**GTCA**GA** | CAGGAGC**C**TC**A**GT**T**G |
| *Oncorhynchus mykiss* | Rainbow trout | 8 | EU524217, EU524221, EU524222, KX145451, LTSM115-12 (BOLD), MOBIL1108-15 (BOLD), MOBIL1110-15 (BOLD), MOBIL1864-16 (BOLD) | Québec, Canada  Ontario, Canada | CC**T**CT**A**GC**C**GGCAAC**C**T | A**AA**TGAAGGGAGAAGAT**A**GT**T**AA**A** | CAGGAGC**C**TC**T**GT**T**G |
| *Oncorhynchus nerka* | Sockeye salmon | 6 | MOBIL1925-16 (BOLD), MOBIL3669-17 (BOLD), MOBIL3692-17 (BOLD), MOBIL4223-17 (BOLD), MOBIL4224-17 (BOLD), WBSF006-15 (BOLD) | Ontario, Canada | CC**T**CTGGC**C**GG**A**AAC**C**T | A**AA**TGAAGGGAGAAGAT**G**GT**T**AAG | C**G**GGAGC**C**TC**T**GT**T**G |
| Oncorhynchus tshawytscha | Chinook salmon | 2 | KX145197, KX145557 | Ontario, Canada | CC**T**CTGGC**C**GGCAAC**C**T | A**AA**TGAAGGGAGAAGAT**C**GTCA**GA** | CAGGAGC**C**TC**A**GT**T**G |
| *Opsopoeodus emiliae* | Pugnose minnow | 1 | OM736889.1 | Pennsylvanie, États-Unis | CC**A**CT**T**GCAGGCAAC**C**T | NA | CAGG**C**GCATC**A**GT**A**G |
| *Osmerus mordax* | Rainbow smelt | 29 | EF609422.1, EU524235.1, EU524236.1, EU524237.1, FJ205604.1, FJ205605.1, FJ205606.1, JQ354253.1, KC015752.1, KC015753.1, KF930207.1, KP257701.1, KP978066.1, KT247719.1, KX145165.1, KX145168.1, KX145169.1, KX145434.1, KX145574.1, MT577162.1, MT577163.1, MT577164.1, MT577165.1, MT577166.1, MW856834.1, MW856836.1, MW856884.1, MW856916.1, OM736824.1 | Manitoba, Canada  Michigan, United States  New-Brunswick, Canada  Newfondland and Labrador, Canada  Ontario, Canada  Sakhalin Oblast, Russia  Lake Superior, United States  Wisconsin, United States  Alaska, United States  Rhode Island, United States | CC**A**CT**T**GC**T**GGCAA**T**TT | AGGTGAAG**A**GAGAA**A**ATTGT**T**AA**A** | C**G**GGAGC**T**TCCGT**A**G |
|  |  |  |  |  |  | AGGTGAAG**A**GA**A**AA**A**ATTGT**T**A**GA** | C**T**GGAGC**T**TC**G**GT**A**G |
|  |  |  |  |  |  |  | CAGGAGC**T**TC**G**GT**A**G |
| *Perca flavescens* | Yellow perch | 11 | EU524240, EU524241, EU524242, EU524243, EU524244, JX516930, KX145007, KX145021, KX145039, KX145260, KX145468 | Québec, Canada  Ontario, Canada | CC**T**CT**T**GC**T**GG**G**AACTT | NA | C**T**GGAGCATC**T**GT**T**G |
| *Percina caprodes* | Logperch | 8 | EU524248, EU524247, EU524246, JN027923, JN027924, KX145151, KX145126, EU524249 | Ontario, Canada  Québec, Canada | CC**TT**T**A**GC**G**GG**A**AACTT | NA | C**C**GG**G**GCATCCGTCG |
| *Percina copelandi* | Channel darter | 9 | EU524251, EU524250, KX145128, KX145420, KX145552, KX145209, KX145044, KX145526, EU524252 | Québec, Canada  Ontario, Canada | CC**T**CTGGC**T**GG**G**AACTT | AG**A**TG**C**AGGGA**A**AAGAT**G**GT**T**AAG | C**C**GGAGCATCCGT**T**G |
| *Percina shumardi* | River darter | 4 | EU524260, KX145208, KX145398, KX145269 | Ontario, Canada | CC**T**CTGGC**C**GG**A**AACTT | NA | C**C**GG**G**GCATCCGT**T**G |
| *Percina maculata* | Blackside darter | 12 | EU524259, EU524258, EU524257, EU524256, EU524255, EU524254, EU524253, KX145112, KX145185, KX145002, KX145062, KX145179 | Ontario, Canada | CCCCTGGC**T**GG**A**AACTT | NA | C**T**GG**G**GCATCCGT**T**G |
| *Percopsis omiscomaycus* | Trout-perch | 13 | EU524261, EU524262, EU524263, EU524264, EU524265, EU524267, EU524268, EU524269, KX145136, KX145144, KX145224, KX145291, KX145311 | Québec, Canada  Ontario, Canada | CCCCTGGC**G**GG**T**AAC**C**T | AGGTG**C**AGGGAGAA**A**ATTGT**G**A**G**G | CAGG**G**GC**C**TCCGTCG |
| *Petromyzon marinus* | Sea lamprey | 10 | EU524270, EU524271, EU524272, EU524273, JN028182, JN028183, JN028184, JN028185, KX145312, KX145569 | Québec, Canada | CCC**T**T**A**GC**C**GG**A**AAC**C**T | NA | C**C**GG**G**GC**C**TC**T**GTCG |
| *Pimephales promelas* | Fathead minnow | 147 | BCFB280-06 (BOLD), EU525085, EU525086, EU525087, EU525088, EU525089, EU525090, EU525091, EU525092, EU525093, EU525094, EU525095, HQ339976, JX516103, JX516109, JX516110, JX516116, JX516119, JX516120, JX516133, JX516134, JX516145, JX516149, JX516154, JX516167, JX516173, JX516191, JX516193, JX516198, JX516206, JX516213, JX516219, JX516229, JX516231, JX516234, JX516252, JX516258, JX516266, JX516268, JX516272, JX516295, JX516299, JX516312, JX516318, JX516322, JX516325, JX516338, JX516372, JX516378, JX516381, JX516391, JX516394, JX516396, JX516397, JX516401, JX516407, JX516418, JX516419, JX516428, JX516431, JX516437, JX516442, JX516444, JX516467, JX516469, JX516479, JX516486, JX516492, JX516493, JX516498, JX516500, JX516504, JX516505, JX516506, JX516508, JX516511, JX516517, JX516519, JX516523, JX516528, JX516530, JX516535, JX516546, JX516548, JX516554, JX516558, JX516569, JX516587, JX516602, JX516609, JX516616, JX516617, JX516622, JX516623, JX516626, JX516629, JX516635, JX516636, JX516665, JX516667, JX516672, JX516677, JX516687, JX516694, JX516696, JX516704, JX516709, JX516714, JX516725, JX516736, JX516737, JX516749, JX516758, JX516760, JX516781, JX516786, JX516807, JX516820, JX516863, JX516892, JX516894, JX516998, JX517076, JX517110, JX517130, JX517142, JX517161, JX517173, JX517194, KT278765.1, KX145000, KX145328, KX145371, KX145431, MG570452.1, MG570454.1, MT577175, MT577176, MT577177, MT577178, NC_028087.1, OK623675.1, OL477723.1, OP418470, OP418471, OP418472, OP418473 | Ontario, Canada  Quebec, Canada,  Manitoba, Canada  Oregon, United States  New York, United States | CC**A**CT**T**GCAGG**T**AA**T**CTT | NA | CAGGAGC**C**TC**A**GT**A**G |
| *Pimephales notatus* | Bluntnose minnow | 81 | EU524276.1, EU525074.1, EU525075.1, EU525076.1, EU525077.1, EU525078.1, EU525079.1, EU525080.1, EU525081.1, EU525082.1, EU525083.1, EU525084.1, HQ557086.1, HQ557087.1, HQ557088.1, HQ557089.1, JN028226.1, JN028227.1, JN028228.1, JN028229.1, JN028230.1, JN028231.1, JN028232.1, JN028233.1, JN028234.1, JN028235.1, JN028236.1, JN028237.1, JX516813.1, JX516825.1, JX516844.1, JX516852.1, JX516860.1, JX516899.1, JX516929.1, JX516983.1, JX517012.1, JX517013.1, JX517016.1, JX517017.1, JX517024.1, JX517025.1, JX517044.1, JX517069.1, JX517075.1, JX517079.1, JX517080.1, JX517094.1, JX517100.1, JX517114.1, JX517116.1, JX517171.1, KF930262.1, KX145117.1, KX145509.1, KX145550.1, MG448640.1, MG448718.1, MG448731.1, MG448808.1, MG448832.1, MG448909.1, MG448933.1, MG449020.1, MG449114.1, MG449528.1, MG449677.1, MG449796.1, MG450276.1, MG450310.1, MG570420.1, MG570450.1, MG570457.1, MG570458.1, MG806797.1, MT455194.1, MT455196.1, MT455557.1, MT456260.1, OM718772.1, OM743017.1 | Ontario, Canada  Québec, Canada  Pennsylvania, United States  Arkansas, United States  Illinois, United States  Kentucky, United States  Maryland, United States  Missouri, United States  Oklahoma, United States  Tennessee, United States  Wisconsin, United States | CC**A**CT**T**GCAGG**T**AAC**C**T |  | CAGGAGCATC**A**GT**A**G |
|  |  |  |  |  | CC**A**CT**T**GCAGG**T**AA**T**CTT |  |  |
|  |  |  |  |  | CC**A**CT**C**GCAGG**T**AA**T**CTT |  |  |
|  |  |  |  |  | CC**A**CT**T**GC**C**GG**T**AA**T**CTT |  |  |
|  |  |  |  |  | CCCCT**ACA**AG**A**C**T**ACTT |  |  |
| *Pomoxis nigromaculatus* | Black crappie | 16 | BIICQ096-18 (BOLD), BIICQ097-18 (BOLD), BIICQ098-18 (BOLD), BIICQ099-18 (BOLD), BIICQ100-18 (BOLD), BIICQ101-18 (BOLD), EU524285, EU524286, EU524287, EU525098, EU525101, EU525102, KX145287, KX145290, KX145449, KX145519 | Québec, Canada  Ontario, Canada | CCC**T**TGGC**C**GGCAAC**C**T | AG**A**TG**G**AGGGAGAAGAT**G**GTCAAG | CAGGAGCATCCGT**T**G |
| *Prosopium cylindraceum* | Round whitefish | 49 | AP013050.1, EU202656.1, EU202657.1, EU524288.1, EU524289.1, EU524290.1, EU524291.1, EU524292.1, EU524293.1, EU524294.1, EU524295.1, EU524296.1, KP978217.1, KP978218.1, KP978219.1, KP978220.1, KP978221.1, KP978222.1, KP978223.1, KP978224.1, KP978225.1, KR733425.1, KT630727.1, KU867893.1, KU867894.1, KU867895.1, KX144996.1, KX145142.1, KX145162.1, KX145501.1, KX145585.1, MF278549.1, MF278550.1, MF278551.1, MF278552.1, MF278553.1, MF278554.1, MF278555.1, MF278556.1, MF278557.1, MF278558.1, MF278559.1, MF278560.1, MF278561.1, MG951550.1, MG951551.1, MT577189.1, MT577190.1, OM736820.1 | Ontario, Canada  Lake Sobachye, Russia  Lake Superior, United States  Alaska, United States  Yukon, Canada | CC**A**CT**A**GCAGGCAAC**C**T | A**A**GTG**T**AGGGAGAAGAT**A**GT**T**AAG | CAGG**G**GC**C**TCCGT**T**G |
|  |  |  |  |  |  | A**A**GTG**T**AGGGA**A**AAGAT**A**GT**T**AAG |  |
|  |  |  |  |  |  | A**A**GTG**G**AGGGAGAAGAT**A**GT**T**AA**A** |  |
| *Pungitius pungitius* | Ninespine stickleback | 14 | EU524321, EU525105, EU525106, EU525107, EU525108, EU525109, EU525110, EU525111, EU525112, KX145024, KX145065, KX145271, KX145302, KX145539 | Québec, Canada  Ontario, Canada | C**A**CCT**A**GCAGG**-**AA**T**TT | AGGTGAAGGGAGAAGATTGT**T**A**G**G | CAGG**T**GC**C**TC**G**GT**T**G |
|  |  |  |  |  |  |  | CAGG**T**GC**C**TC**A**GT**T**G |
| *Rhinichthys cataractae* | Longnose dace | 44 | MG421536.1, MG421490.1, MG421091.1, MG421079.1, MG421051.1, MG423205.1, MG423156.1, MG423028.1, MG422434.1, MG422400.1, MG422246.1, KX145530.1, KX145316.1, KX145220.1, KX145140.1, KX145137.1, JN028362.1, JN028360.1, JN028361.1, EU525130.1, EU525129.1, EU525128.1, EU525127.1, EU525126.1, EU525125.1, EU525124.1, EU525123.1, EU525122.1, EU525121.1, EU524327.1, EU524326.1, EU524325.1, EU524324.1, EU524323.1, MT667247.1, MG570446.1, MG570448.1, OL404932.1, OM736844.1, OR031097.1, OR031098.1, OK336456.1, OK623674.1, OL693869.1 | Manitoba, Canada  Québec, Canada  Ontario, Canada  Pennsylvania, United States  New York, United States  Oregon, United States | CC**G**CT**C**GCAGGCAAC**C**T | AGGTG**T**A**AA**GAGAAGATTGT**G**A**G**G | CAGGAGCATC**A**GT**A**G |
|  |  |  |  |  |  | AGGTG**T**AG**A**GAGAAGAT**C**GT**G**A**G**G |  |
| *Rhinichthys atratulus* | Eastern blacknose dace | 18 | PP958238.1, MG570445.1, MG570444.1, JX516971.1, JX516988.1, JX516968.1, JX516962.1, JX516960.1, JX516926.1, JX516914.1, JX516905.1, EU525120.1, EU525119.1, EU525118.1, EU525117.1, EU525116.1, EU525115.1, EU524322.1 | Connecticut, United States  New York, United States  Québec, Canada  New-Brunswick, Canada | CC**G**CT**C**GCAGG**T**AA**CC**T | AGGTG**T**AG**A**GAGAAGATTGT**G**A**G**G | CAGGAGCATC**A**GT**A**G |
| *Rhinichthys obtusus* | Western blacknose dace | 103 | MG570417.1, NC_037010.1, JX517192.1, JX517191.1, JX517190.1, JX517187.1, JX517184.1, JX517182.1, JX517180.1, JX517176.1, JX517167.1, JX517156.1, JX517155.1, JX517152.1, JX517151.1, JX517144.1, JX517143.1, JX517141.1, JX517133.1, JX517137.1, JX517131.1, JX517127.1, JX517118.1, JX517115.1, JX517112.1, JX517107.1, JX517096.1, JX517089.1, JX517088.1, JX517087.1, JX517086.1, JX517077.1, JX517068.1, JX517062.1, JX517060.1, JX517058.1, JX517043.1, JX517041.1, JX517034.1, JX517031.1, JX517028.1, JX517022.1, JX517019.1, JX517015.1, JX517006.1, JX517004.1, JX517003.1, JX517000.1, JX516980.1, JX516973.1, JX516972.1, JX516970.1, JX516967.1, JX516963.1, JX516956.1, JX516953.1, JX516947.1, JX516942.1, JX516939.1, JX516936.1, JX516935.1, JX516928.1, JX516919.1, JX516912.1, JX516911.1, JX516903.1, JX516902.1, JX516900.1, JX516896.1, JX516895.1, JX516891.1, JX516890.1, JX516886.1, JX516884.1, JX516883.1, JX516882.1, JX516880.1, JX516875.1, JX516868.1, JX516861.1, JX516856.1, JX516850.1, JX516839.1, JX516838.1, JX516837.1, JX516835.1, JX516834.1, JX516830.1, JX516823.1, JX516822.1, JX516819.1, JX516809.1, JX516802.1, JX516801.1, JX516799.1, EU525134.1, EU525133.1, EU525132.1, EU525131.1, EU524336.1, EU524335.1, EU524334.1, EU524333.1 | Wisconsin, United States  Illinois, United States  Ontario, Canada | CC**A**CT**C**GCAGG**T**AAC**C**T | AGGTGAAG**A**GAGAAGATTGT**G**A**G**G | CAGGAGCATC**A**GT**A**G |
| *Salmo salar* | Atlantic salmon | 15 | EMRKT067-07 (BOLD), EMRKT088-07 (BOLD), EU524349, EU524351, EU524352, EU524353, LTSM108-12 (BOLD), LTSM139-12 (BOLD), LTSM145-12 (BOLD), LTSM205-12 (BOLD), LTSM260-12 (BOLD), LTSM308-12 (BOLD), LTSM352-12 (BOLD), LTSM381-12 (BOLD), LTSM393-12 (BOLD) | Ontario, Canada | CC**T**CT**A**GCAGG**T**AA**T**CTT | NA | CAGGAGC**T**TCCGT**T**G |
| *Salmo trutta* | Brown trout | 19 | EU524354, EU524355, EU524356, KX145471, KX145518, MOBIL10206-19 (BOLD), MOBIL10208-19 (BOLD), MOBIL10209-19 (BOLD), MOBIL10231-19 (BOLD), MOBIL10232-19 (BOLD), MOBIL10297-19 (BOLD), MOBIL10335-19 (BOLD), MOBIL10339-19 (BOLD), MOBIL10340-19 (BOLD), MOBIL10343-19 (BOLD), MOBIL10364-19 (BOLD), MOBIL10731-20 (BOLD), MOBIL10749-20 (BOLD), MOBIL10769-20 (BOLD) | New-Brunswick, Canada  Alberta, Canada | CC**T**CT**A**GC**C**GGCAA**T**CTT | NA | CAGGAGC**T**TCCGT**T**G |
| *Salvelinus alpinus* | Arctic char | 10 | ETOHS001-06 (BOLD), ETOHS002-06 (BOLD), EU524357, EU524358, EU524359, EU524360, EU524361, EU524362, EU524363, KY122048 | Québec, Canada | CC**T**CT**A**GC**C**GG**G**AAC**C**T | A**AA**TGAAGGGAGAAGAT**A**GT**T**AA**A** | CAGG**G**GC**C**TCCGT**T**G |
| *Salvelinus fontinalis* | Brook trout | 12 | ETOHS014-06 (BOLD), EU522404, EU522405, EU522406, EU522407, EU522408, EU522409, EU524366, EU524367, KX145042, KX145127, KX145572 | Québec, Canada | CC**T**CT**A**GC**T**GG**G**AAC**C**T | NA | CAGGAGC**T**TCCGT**T**G |
| *Salvelinus namaycush* | Lake trout | 9 | ETOHS009-06 (BOLD), ETOHS010-06 (BOLD), EU522422, EU522423, EU522424, EU522425, KX145143, KX145531, KX145595 | Québec, Canada  Ontario, Canada | CC**T**CT**A**GC**C**GG**G**AAC**C**T | NA | CAGG**G**GC**C**TCCGT**T**G |
| *Sander canadensis* | Sauger | 6 | EU524368, EU524369, EU524370, EU524371, EU524372, EU524373 | Québec, Canada | NA | NA | C**C**GG**G**GCATCCGT**T**G |
| *Sander vitreus* | Walleye | 44 | EU524374, EU524375, EU524376, EU524377, EU524378, EU524379, EU524380, KP978283, KP978284, KP978285, KP978286, KP978287, KP978288, KP978290, KP978291, KP978292, KP978293, KP978294, KP978295, KP978296, KP978297, KP978298, KP978299, KP978300, KP978301, KP978302, KP978303, KP978304, KP978305, KP978306, KP978307, KP978308, KP978310, KP978312, KX145106, KX145115, KX145158, KX145180, KX145279, MNRFE001-14 (BOLD), MNRFE016-14 (BOLD), MNRFE017-14 (BOLD), MNRFE018-14 (BOLD), MOBIL1076-15 (BOLD) | Québec, Canada  Ontario, Canada | NA | NA | C**C**GG**G**GCATC**T**GTCG |
| *Semotilus atromaculatus* | Creek chub | 15 | EU525136, EU525137, EU525138, EU525139, EU525140, EU525141, EU525142, EU525143, EU525144, HQ557726, KX144980, KX145214, KX145414, KX145426, KX145494 | Québec, Canada  Ontario, Canada | CC**A**CT**T**GC**G**GG**T**AA**T**CTT | NA | C**C**GGAGCATC**A**GT**A**G |
| *Semotilus corporalis* | Fallfish | 12 | EU524383, EU525145, EU525146, EU525147, EU525148, EU525149, EU525150, EU525151, EU525152, KX145325, KX145500, KX145605 | Québec, Canada  Ontario, Canada | NA | AGGTG**C**AG**A**GAGAAGATTGTCA**G**G | C**C**GG**G**GCATCCGT**T**G |
| *Tinca tinca* | Tench | 7 | EU525154, EU525157, EU525158, EU525159, EU525160, EU525161, EU525162 | Québec, Canada | CC**A**CT**C**GCAGG**T**AA**T**CTT | AGGTGAAG**T**GAGAA**A**ATTGT**T**A**G**G | CAGGAGC**C**TC**A**GT**A**G |
| *Umbra limi* | Central mudminnow | 14 | EU522446, EU522447, EU522448, EU522449, EU522450, EU522451, EU522452, EU522453, EU524391, KX145103, KX145123, KX145277, KX145361, KX145415 | Québec, Canada  Ontario, Canada | CCCCTGGC**T**GGCAAC**C**T | AGGTG**G**AGGGA**A**AAGAT**A**GT**T**A**G**G | C**C**GG**C**GC**C**TCCGT**A**G |

2) In vitro validation and assay sensitivity
a) Preliminary primer screening was performed with FAST SYBR Green (Life Technologies, Carlsbad, CA). Amplifications were performed on a 7,500 Fast Real-Time PCR System (Applied Biosystems, Waltham, MA) in a final volume of 20 μL: 10 μL of Fast SYBR® Green Master Mix (SYBR, LLC, Hanover, PA), 1 μL of each primer (10 μM), 2 μL of DNA (5–10 ng) and 6 μL of UltraPure Distilled Water (DNAse, RNAse, Free, Invitrogen^TM^, Waltham, MA) following these conditions: 95°C for 20 s, 40 cycles × [95°C for 3 s, 60°C for 30 s]. Finally, selected primers were tested with their probes in a TaqMan (Applied Biosystems) assay in a final volume of 20 μL including 0.4 μL of primer F (10 μM), 1.2 μL of primer R (10 μM), 0.4 μL of probe (10 μM), 10 μL of TaqPath ProAmp (2x)®(Applied Biosystems), BSA 0.4mg/mL, 4.2 μL of dH2O and 3 μL of DNA (10 ng) following these conditions: 50 °C for 2 min, 95 °C for 10 min 50 cycles × [95 °C for 15 s, 63 °C for 1 min]. Amplifications were performed on a QuantStudio™ 3 (Applied Biosystems).
b) *Specificity results*

A standard curve was performed following the same conditions as described above for the TaqMan assay. A reference plasmid containing the target amplicon sequence was designed from the COI gene sequence. A dilution series was prepared in a sterile yeast tRNA (100 μg/μL) solution. Concentrations and number of replicates used for the standard curve are provided in Table S3.

Table S3. Properties of the standard curve to determine the limit of detection and limit of quantification of round goby DNA. The concentration of the initial plasmid DNA solution used for the dilution series was determined using a QuantStudio AbsoluteQ Digital PCR System (Thermo Fisher Scientific, Waltham, MA, USA).

| Concentration (copies/reaction) | Replicates |
| --- | --- |
| 6250 | 6 |
| 1250 | 6 |
| 250 | 6 |
| 50 | 6 |
| 10 | 6 |
| 2 | 16 |
| 0.4 | 16 |
| 0.08 | 16 |
| 0 | 16 |

The data relating to the calibration curve and its linear regression were modeled according to the R script (R Core Team, 2025) from Klymus et al., 2020 (Table S4).

The data relating to the limit of detection (LOD) and the limit of quantification (LOQ) were modeled according to the R eLowQuant script of Lesperance et al. 2021.

**Table S4.** Characteristics of the qPCR assay sensitivity. R^2^ = regression coefficient of the standard curve, LOD = limit of detection, LOQ = limit of quantification.

| **Linear regression slope** | **y-intercept** | **R^2^** | **Amplification efficiency (%)** | **LOD (copies/reaction)** | **LOQ (copies/reaction)** |
| --- | --- | --- | --- | --- | --- |
| -3.3358 | 37.6357 | 0.995 | 94.423 | 0.3 | 1 |

c) *Species for cross-amplification tests*

All species used for cross-amplification tests are presented in Table S5. No cross-amplification with the DNA of all related tested species was detected.

**Table S5.** Information on the provenance and amplification of the target species (*Neogobius melanostomus*) and non-target species. The DNA concentration used for all non-target species tested was 0.01 ng/μL.

| **Scientific name** | **Common name** | **ID** | **Origin** | **Amplification** |
| --- | --- | --- | --- | --- |
| *Neogobius melanostomus* | Round goby | S_23_00737 | Québec, Canada | Positive |
| *Neogobius melanostomus* | Round goby | S_23_00767 | Québec, Canada | Positive |
| *Neogobius melanostomus* | Round goby | S_23_00787 | Québec, Canada | Positive |
| *Neogobius melanostomus* | Round goby | S_25_00194 | Québec, Canada | Positive |
| *Neogobius melanostomus* | Round goby | S_25_00195 | Québec, Canada | Positive |
| *Neogobius melanostomus* | Round goby | S_25_00196 | Québec, Canada | Positive |
| *Percina copelandi* | Channel darter | S_24_09242 | Québec, Canada | Negative |
| *Percina copelandi* | Channel darter | S_24_09154 | Ontario, Canada | Negative |
| *Percina caprodes* | Logperch | S_24_09234 | Ontario, Canada | Negative |
| *Percina caprodes* | Logperch | S_24_09235 | Ontario, Canada | Negative |
| *Percina caprodes* | Logperch | S_24_09146 | Ontario, Canada | Negative |
| *Percina caprodes* | Logperch | S_24_09147 | Québec, Canada | Negative |
| *Ammocrypta pellucida* | Eastern sand darter | S_24_09226 | Ontario, Canada | Negative |
| *Ammocrypta pellucida* | Eastern sand darter | S_24_09227 | Ontario, Canada | Negative |
| *Ammocrypta pellucida* | Eastern sand darter | S_24_09228 | Ontario, Canada | Negative |
| *Notropis volucellus* | Mimic shiner | S_24_03748 | Lac Saint-François, Canada | Negative |
| *Notropis volucellus* | Mimic shiner | S_24_03801 | Québec, Canada | Negative |
| *Notropis volucellus* | Mimic shiner | S_24_03802 | Québec, Canada | Negative |
| *Notropis stramineus* | Sand shiner | S_24_03785 | Québec, Canada | Negative |
| *Notropis stramineus* | Sand shiner | S_24_03789 | Québec, Canada | Negative |
| *Notropis stramineus* | Sand shiner | S_24_03792 | Ontario, Canada | Negative |
| *Notropis rubellus* | Rosyface shiner | S_24_03751 | Québec, Canada | Negative |
| *Notropis rubellus* | Rosyface shiner | S_24_03815 | Québec, Canada | Negative |
| *Notropis hudsonius* | Spottail shiner | S_24_03774 | Québec, Canada | Negative |
| *Notropis hudsonius* | Spottail shiner | S_24_03775 | Québec, Canada | Negative |
| *Notropis hudsonius* | Spottail shiner | S_24_03776 | Québec, Canada | Negative |
| *Notropis heterodon* | Blackchin shiner | S_24_03747 | Québec, Canada | Negative |
| *Notropis heterodon* | Blackchin shiner | S_24_03758 | Québec, Canada | Negative |
| *Notropis heteroplesis* | Blackchin shiner | S_24_03766 | Québec, Canada | Negative |
| *Notropis heteroplesis* | Blackchin shiner | S_24_03767 | Québec, Canada | Negative |
| *Notropis heteroplesis* | Blackchin shiner | S_24_03769 | Québec, Canada | Negative |
| *Notropis buchanani* | Ghost shiner | S_24_09210 | Ontario, Canada | Negative |
| *Notropis bifrenatus* | Bridle shiner | S_24_03778 | Québec, Canada | Negative |
| *Notropis bifrenatus* | Bridle shiner | S_24_03779 | Québec, Canada | Negative |
| *Notropis bifrenatus* | Bridle shiner | S_24_03783 | Québec, Canada | Negative |
| *Notropis anogenus* | Pugnose shiner | S_24_03749 | Ontario, Canada | Negative |
| *Notropis anogenus* | Pugnose shiner | S_24_09125 | Ontario, Canada | Negative |
| *Notropis anogenus* | Pugnose shiner | S_24_09127 | Ontario, Canada | Negative |
| *Notropis atherinoides* | Emerald shiner | S_24_03821 | Québec, Canada | Negative |
| *Notropis atherinoides* | Emerald shiner | S_24_03822 | Québec, Canada | Negative |
| *Notropis atherinoides* | Emerald shiner | S_24_03823 | Québec, Canada | Negative |
| *Myoxocephalus thompsonii* | Deepwater sculpin | S_24_09345 | Ontario, Canada | Negative |
| *Myoxocephalus thompsonii* | Deepwater sculpin | S_24_09346 | Ontario, Canada | Negative |
| *Myoxocephalus thompsonii* | Deepwater sculpin | S_24_09193 | Ontario, Canada | Negative |
| *Myoxocephalus quadricornis* | Fourhorn sculpin | S_24_09182 | Ontario, Canada | Negative |
| *Myoxocephalus quadricornis* | Fourhorn sculpin | S_24_09183 | Ontario, Canada | Negative |
| *Hybognathus hankinsoni* | Brassy minnow | S_24_09356 | Ontario, Canada | Negative |
| *Hybognathus regius* | Eastern silvery minnow | S_24_09351 | Québec, Canada | Negative |
| *Hybognathus regius* | Eastern silvery minnow | S_24_09354 | Québec, Canada | Negative |
| *Cottus bairdii* | Mottled sculpin | S_24_09134 | Québec, Canada | Negative |
| *Cottus bairdii* | Mottled sculpin | S_24_09135 | Ontario, Canada | Negative |
| *Cottus bairdii* | Mottled sculpin | S_24_09193 | Ontario, Canada | Negative |
| *Cottus cognatus* | Slimy sculpin | S_23_01699 | Québec, Canada | Negative |
| *Cottus cognatus* | Slimy sculpin | S_23_01700 | Québec, Canada | Negative |
| *Cottus cognatus* | Slimy sculpin | S_23_01702 | Québec, Canada | Negative |
| *Cottus ricei* | Spoonhead sculpin | S_23_01701 | Québec, Canada | Negative |
| *Cottus ricei* | Spoonhead sculpin | S_23_01706 | Québec, Canada | Negative |
| *Cottus ricei* | Spoonhead sculpin | S_23_01707 | Québec, Canada | Negative |
| *Etheostoma caeruleum* | Rainbow darter | S_24_09211 | Ontario, Canada | Negative |
| *Etheostoma caeruleum* | Rainbow darter | S_24_09212 | Ontario, Canada | Negative |
| *Etheostoma caeruleum* | Rainbow darter | S_24_09213 | Ontario, Canada | Negative |
| *Etheostoma flabellare* | Fantail darter | S_24_09236 | Québec, Canada | Negative |
| *Etheostoma flabellare* | Fantail darter | S_24_09238 | Québec, Canada | Negative |
| *Etheostoma flabellare* | Fantail darter | S_24_09239 | Québec, Canada | Negative |
| *Etheostoma nigrum* | Johnny darter | S_24_09221 | Québec, Canada | Negative |
| *Etheostoma nigrum* | Johnny darter | S_24_09223 | Québec, Canada | Negative |
| *Etheostoma nigrum* | Johnny darter | S_24_09225 | Québec, Canada | Negative |
| *Etheostoma olmstedi* | Tessellated darter | S_24_09217 | Québec, Canada | Negative |
| *Etheostoma olmstedi* | Tessellated darter | S_24_09218 | Québec, Canada | Negative |
| *Etheostoma olmstedi* | Tessellated darter | S_24_09220 | Québec, Canada | Negative |
| *Exoglossum maxillingua* | Cutlips minnow | S_24_09578 | Québec, Canada | Negative |
| *Exoglossum maxillingua* | Cutlips minnow | S_24_09579 | Québec, Canada | Negative |
| *Exoglossum maxillingua* | Cutlips minnow | S_24_09580 | Québec, Canada | Negative |
| *Rhinichthys atratulus* | Eastern blacknose dace | S_24_03818 | Québec, Canada | Negative |
| *Rhinichthys atratulus* | Eastern blacknose dace | S_24_03819 | Québec, Canada | Negative |
| *Rhinichthys atratulus* | Eastern blacknose dace | S_24_03820 | Québec, Canada | Negative |
| *Rhinichthys obtusus* | Western blacknose dace | S_24_09132 | Ontario, Canada | Negative |
| *Rhinichthys obtusus* | Western blacknose dace | S_24_09133 | Ontario, Canada | Negative |
| *Rhinichthys cataractae* | Longnose dace | S_24_03829 | Québec, Canada | Negative |
| *Rhinichthys cataractae* | Longnose dace | S_24_03832 | Québec, Canada | Negative |
| *Rhinichthys cataractae* | Longnose dace | S_24_03833 | Québec, Canada | Negative |
| *Pimephales promelas* | Fathead minnow | S_24_03753 | Québec, Canada | Negative |
| *Pimephales promelas* | Fathead minnow | S_24_03754 | Québec, Canada | Negative |
| *Pimephales promelas* | Fathead minnow | S_24_03756 | Québec, Canada | Negative |
| *Pimephales notatus* | Bluntnose minnow | S_24_03806 | Québec, Canada | Negative |
| *Pimephales notatus* | Bluntnose minnow | S_24_03807 | Québec, Canada | Negative |
| *Pimephales notatus* | Bluntnose minnow | S_24_03808 | Québec, Canada | Negative |
| *Luxilus cornutus* | Common shiner | S_24_03825 | Québec, Canada | Negative |
| *Luxilus cornutus* | Common shiner | S_24_03827 | Québec, Canada | Negative |
| *Luxilus cornutus* | Common shiner | S_24_03828 | Québec, Canada | Negative |
| *Cyprinella spiloptera* | Spotfin shiner | S_24_03811 | Québec, Canada | Negative |
| *Cyprinella spiloptera* | Spotfin shiner | S_24_03813 | Québec, Canada | Negative |
| *Cyprinella spiloptera* | Spotfin shiner | S_24_03814 | Québec, Canada | Negative |
